# How much is ‘enough’? Considerations for functional connectivity reliability in pediatric naturalistic fMRI

**DOI:** 10.1101/2024.12.02.626468

**Authors:** Shefali Rai, Kate J. Godfrey, Kirk Graff, Ryann Tansey, Daria Merrikh, Shelly Yin, Matthew Feigelis, Damion V. Demeter, Tamara Vanderwal, Deanna J. Greene, Signe Bray

## Abstract

functional connectivity (FC) measurements are important for robust and reproducible findings, yet pediatric functional magnetic resonance imaging (fMRI) faces unique challenges due to head motion and bias toward shorter scans. Passive viewing conditions during fMRI offer advantages for scanning pediatric populations, but FC reliability under these conditions remains underexplored. Here, we used precision fMRI data collected across three passive viewing conditions to directly compare FC reliability profiles between 25 pre-adolescent children and 25 adults, with each participant providing over 2.8 hours of data over four sessions. We found that FC test-retest correlations increased asymptotically with scan length, with children requiring nearly twice the post-censored scan time (24.6 minutes) compared to adults (14.4 minutes) to achieve comparable reliability, and that this effect was only partly attributable to head motion. Reliability differences between lower-motion adults and higher-motion children were spatially non-uniform and largest in ventral anterior temporal and frontal regions. While averaging features within functional networks improved intraclass correlation coefficient (ICC) reliability, values for higher-motion children remained in the poor-to-fair ICC range even with 48 minutes of total scan time. Viewing conditions with greater engagement reduced head motion in children but had lower FC reliability compared to less engaging ‘low-demand’ videos, suggesting complex state- or condition-related trade-offs. These findings have important implications for developmental neuroimaging study design, particularly for higher motion pediatric populations.

## 1 Introduction

Reliable measurements – defined in this context as measures with high test-retest consistency and minimal variation across repeated assessments (Shrout & Fleiss, 1979) – are necessary for reproducible research findings. While a growing body of literature supports best practices for reliable functional magnetic resonance imaging functional connectivity (fMRI-FC) in adults (Bennett & Miller, 2010, 2013; Elliott et al., 2021; Kragel et al., 2021; Noble et al., 2019a; Shehzad et al., 2009), and despite research on both clinical and nonclinical pediatric FC reliability (Camp et al., 2024; Somandepalli et al., 2015; Thomason et al., 2011; Y. Wang et al., 2021), there are few established guidelines specific to pediatric fMRI-FC reliability. Relatedly, precision fMRI (pfMRI) is an emerging neuroimaging approach that can reliably estimate metrics of brain function from individuals by collecting relatively large amounts of fMRI data per person (Michon et al., 2022). Adult studies using pfMRI describe asymptotic relationships between scan time and FC reliability (Gordon et al., 2017; Laumann et al., 2015), but the shape of these curves has not been compared between adult and pediatric populations, which limits our understanding of its applicability to children and other populations with substantial head motion. Understanding how FC reliability profiles differ between adults and children and how factors such as scan length, head motion, and viewing conditions impact these profiles, is critical for advancing reproducible developmental neuroimaging.

Reliability of FC measures in pediatric populations remains a significant challenge due to head motion and concerns regarding tolerability of longer scans (Dosenbach et al., 2017). Some studies also suggest that lower FC reliability in developmental populations may be inherent to developmental differences in functional brain organization (i.e., higher intra-individual variability), and not just a reflection of noise (Bottenhorn et al., 2023; Vanderwal et al., 2021). In adult populations, prior work has suggested that whole functional connectome reliability asymptotes between 30 – 90 minutes of data, depending on factors including scan sequence and level of resolution (Gordon et al., 2017; Noble, Spann, et al., 2017; Noble et al., 2019a; Shah et al., 2016); though, these estimates may not be applicable for higher motion pediatric cohorts. Previous research has highlighted that both age and motion affect FC reliability (Parkes et al., 2018; Shirer et al., 2015; Song et al., 2012); however, to our knowledge, no studies have collected longer duration, multi-session fMRI time series to directly compare FC time-by-reliability profiles between young children and adults, while also assessing the potential impacts of head motion on the reliability of post-censored data.

Given the need to reduce head motion and improve participant engagement for reliable FC measures, movies offer a compelling alternative to both conventional tasks requiring focused attention and task-free rest (Sonkusare et al., 2019; Vanderwal et al., 2019). Movie-watching fMRI has proven valuable for populations that are difficult to scan, such as young children, due to concerns with task performance, higher head motion, drowsiness, and scanner-related anxiety (Cantlon & Li, 2013; Eickhoff et al., 2020; Engelhardt et al., 2017; Frew et al., 2022; Greene et al., 2018; Vanderwal et al., 2015).

While conventional tasks variably influence FC reliability (Elliott et al., 2019; Rai et al., 2024; Shah et al., 2016), naturalistic conditions have demonstrated comparable or superior FC reliability compared to both resting-state and conventional task conditions (Elliott et al., 2019; Hasson et al., 2010; O’Connor et al., 2017; Vanderwal et al., 2021; J. Wang et al., 2017a). Recent reviews (Eickhoff et al., 2020; Nastase et al., 2020; Sonkusare et al., 2019) further highlight the benefits of naturalistic paradigms for developmental neuroimaging, emphasizing increased compliance and ecological validity. However, there remains limited guidance on how choice of video may impact FC reliability for children and youth.

Canonical functional brain networks have been shown to exhibit varying levels of FC reliability, with higher-order cognitive networks (e.g., default mode, frontoparietal and attention networks) generally demonstrating greater reliability relative to sensory networks (e.g., somatomotor and visual networks) (Noble, Spann, et al., 2017; Rai et al., 2024; Tozzi et al., 2020). Investigating the variation in reliability patterns and their interaction with head motion is important for planning analyses in pediatric fMRI studies, especially for higher-order cognitive networks that are often of interest in developmental and clinical research (Grayson & Fair, 2017; Marek et al., 2015; Uddin et al., 2009). Hence, for this study, we considered variation across functional networks in the relationship between scan duration and reliability.

This study used a densely sampled dataset from adults and pre-adolescent children (aged 6.6 – 8.9 years) to compare time-by-reliability relationships while considering age, head motion, brain networks and viewing condition. While FC test-retest reliability is expected to be most reliable between identical repeated sessions (e.g., having participants watch the same movie scene at each measurement) (O’Connor et al., 2017; J. Wang et al., 2017a), here we consider reliability across different movies with similar content (e.g., different scenes from the same narrative film). Recent findings have demonstrated that ICC measures remain consistent across different movies (Tian et al., 2021) and have found reliable functional network topography using different films (Jiahui et al., 2023).

First, we compared test-retest correlation profiles between adults and children with increasing scan lengths while considering the impact of head motion. Next, we examined the spatial distribution of FC reliability across the cortex in high- and low-motion groups using both test-retest correlation and ICC. Finally, we assessed the effect of different passive viewing conditions on data retention and test-retest correlation. Through this work, we provide some practical guidelines for developmental fMRI researchers, particularly when working with higher motion pediatric populations, with the limitation that data acquisition, processing choices and sample characteristics can all impact the specific values reported here.

## 2 Methods

### 2.1 Participants

Data were collected from 50 participants consisting of 25 parent-child pairs (5 female-female, 7 female-male, 8 male-female, 5 male-male), recruited through community advertisements and a laboratory participant database. Children were typically developing between the ages of 6.56 - 8.92 years old at session 1, (mean – 7.88 years, std – 0.69 years; 13 F, 12 M), and parents were between the ages of 33.75 - 47.13 years old at session 1 (mean – 41.39 years, std – 3.63 years; 12 F, 13 M). One child participant and their parent were excluded from analysis due to excessive motion (< 40 minutes of post-censored data), resulting in a final sample of 24 parent-child pairs. Our sample included two families in which two parents and two children participated. Children provided assent, while parents provided written informed consent for their own and their child’s participation in the study. Study procedures were approved by the University of Calgary Conjoint Health Research Ethics Board (REB19-1973).

### 2.2 MRI data collection

Participants completed four MRI sessions in a 3T GE MR750w (Waukesha, WI) scanner with a 32-channel head coil, approximately one week apart, at the Alberta Children’s Hospital. EEG and cognitive data were also collected as part of this study which are not reported here. The order was pseudo-randomized, with half the parents undergoing EEG first, followed by MRI, while their children had MRI first, then EEG, and half the sample in a reversed order. In each MRI session, a T1-weighted 3D BRAVO sequence was acquired (0.8 mm isotropic, TR = 6.764 ms, TE = 2.908 ms, FA = 10°). Multi-echo, multiband functional data were collected in six runs per session using a T2*-weighted gradient-echo echo-planar sequence covering the full brain (3.4 mm in-plane, 45 slices, slice thickness = 3mm, slice spacing=0.4mm, TR = 2 s, TEs = 13, 32.3, 51.6 ms, FA = 70°, FOV = 220 mm, AP phase encoding direction, phase acceleration=2, multiband acceleration factor = 3). A total of 205 volumes (excluding 5 dummy volumes) were acquired per run, for a total acquisition time of roughly 41 minutes per session and approximately 2.8 hours per participant across 4 sessions.

In each session, functional data were collected in two runs for each of three passive viewing conditions. Participants watched new content in all conditions and at all scan sessions (i.e., movie scenes, clips, and videos were not repeated):

#### Narrative Movie

Sequential scenes from the live-action movie *Dora and the Lost City of Gold* (2019). This movie was selected based on the following criteria: (1) lack of visually dark and scary scenes appropriate for viewing on a projector in the MRI scanner and for our pre-adolescent sample; (2) filmed within the last decade; (3) highly engaging Hollywood live-action movie (Cutting et al., 2011), and (4) relatively diverse cast.

#### Non-Narrative Clips

Short (24-65 seconds long, mean = 46.4 sec) and popular (> 1 million views at the time of download) videos accessed via *YouTube* or *TikTok* were selected to sustain visual interest with minimal storytelling (e.g., video game scenes such as *Minecraft*, stop-motion animation, miniature food preparation videos a.k.a. ‘tiny kitchen’, toy unboxing videos, *TikTok* dancing videos, Rube Goldberg machines, and building videos where individuals create structures such as water slides using natural materials). Background music was either kept from the original clips or instrumental music was added to clips without sound.

#### Low-Demand Video

In pilot testing, participants reported sleepiness during Inscapes (Vanderwal et al., 2015) across repeated viewings. We therefore developed a condition consisting of calming videos inspired by Inscapes but with more variety across sessions. For this condition, videos featured gentle instrumental music and slow-moving imagery (e.g., Earth seen from space, underwater wildlife footage, walk through a canyon, and drone footage of a Swiss farmland or a Japanese island).

### 2.3 Behavioural data collection

Following each MRI session, participants completed surveys (Supplemental Table 1) that asked recall questions about the videos they viewed during each of 6 functional runs and whether they felt tired, sleepy, or had difficulty staying awake (derived from the PROMIS Sleep-Related Impairment Short Form (Yu et al., 2012)). To assess attention to videos, the percent of correct responses were calculated. To assess drowsiness levels, a summary score was computed by coding no response as 0, felt tired as 1, sleepy as 2, difficulty staying awake as 3, which were summed across the two runs per viewing condition.

### 2.4 fMRI Preprocessing

We used a customized preprocessing pipeline in Nipype version 1.5.0 (Gorgolewski et al., 2011) with tools from FSL version 6.0.0 (Smith et al., 2004), ANTs version 2.3.4 (Avants et al., 2011), and AFNI version 21.1.16 (Cox, 1996) to preprocess individual scans.

For T1-weighted image processing, ANTs was used to correct intensity inhomogeneities, remove the skull and non-brain tissues, warp each participant’s brain to either the adult MNI 152 nonlinear atlas or the pediatric NIHPD 7–11-year-old atlas (Fonov et al., 2009, 2011), generate tissue segmentations, and then warp segmentations back into native space. Following this, FSL was used to create white matter, cerebrospinal fluid, and grey matter masks and AFNI was used to erode tissue segments by 7 levels.

Initial preprocessing steps were done on each echo separately. The first 5 volumes for each run were discarded using FSL *ExtractROI*. Using FSL *MCFLIRT*, multi-echo EPI (MEPI) images were warped to the first volume of each run, to generate motion estimates on the uncorrected data, and for rigid body realignment. Framewise displacement (FD) estimates from the second echo of each run were used as motion estimates going forward. FSL *slicetimer* was used for slice time correction. Next, FSL *FLIRT* was used to co-register echoes two and three to the first echo, as a reference, for each video run separately, i.e., for session 1, narrative run 1, echo 1 is used as the reference image to co-register session 1 narrative run 1 echo 2 and echo 3. Finally, FSL *Merge* was used to combine all three echoes for each video run together. Next, FSL *FLIRT* was used to co-register all videos for each session. Using session 1 echo 1 as the reference, the remaining sessions and echoes were co-registered to that reference image. Finally, we used FSL *MERGE* to concatenate all six runs from all conditions together.

### 2.4 MEPI Denoising and Optimal Combination

For each session, the merged 3-echo inputs were denoised and optimally combined using the *tedana* workflow version 0.0.12, a Python library to process multi-echo fMRI data (DuPre et al., 2021). Multi-echo data were optimally combined using the T2* combination method (Posse et al., 1999). Among the three options for selecting principal component analysis (PCA) components, our study opted for the Akaike Information Criterion option. This choice, known as the least aggressive approach, improves the probability of successful convergence in the following independent component analysis (ICA) step, while retaining meaningful components. ICA was then used to classify components as BOLD (TE-dependent), non-BOLD (TE-independent), or uncertain (low-variance) based on the Kundu decision tree (v2.5) (Kundu et al., 2013). ICA-based denoising methods tend to fail at classifying many of the components labelled as uncertain (low-variance components) (Dipasquale et al., 2017; Griffanti et al., 2017; Mejia et al., 2020); therefore, we reviewed and manually re-classified components as necessary.

### 2.5 Optimally Combined MEPI Processing

Subsequent steps were run on individual optimally combined (OC) fMRI runs (4 sessions x 3 viewing conditions x 2 runs). FSL *BET* was used for skull stripping, and FSL *FLIRT* was used for boundary-based registration to register each OC image to the participants’ T1-weighted image. Next, linear regression was performed to remove the mean, linear, and quadratic trends from each voxel, along with bandpass temporal filtering between 0.01 – 0.08 Hz, and nuisance regression. We filtered high-frequency motion (>0.1 Hz) in the phase-encoding direction (Gratton, Dworetsky, et al., 2020) from head motion estimates, and following that, regressed out 24 head motion parameters, along with white matter, cerebrospinal fluid, and the global signal. Lastly, we censored volumes above a FD threshold of 0.15 mm (Power et al., 2012); this threshold was chosen after inspection of the FD ’floor’ across participants which showed that for low-motion participants essentially all volumes fell below this threshold. Preprocessed functional runs were then warped to the participants’ T1 using ANTS *ApplyTransforms* and runs of each viewing condition were concatenated.

### 2.6 Cortical surface generation

T1-weighted images were used to generate the cortical surface of each participant through Freesurfer’s *recon-all* pipeline (Dale et al., 1999), version 6.0. Subsequently, the outputs of Freesurfer were converted into grayordinate-based space in the Connectivity Informatics Technology Initiative (CIFTI) format (Glasser et al., 2013a) using Ciftify’s *ciftify_recon_all* pipeline (Dickie et al., 2019).The grayordinate resolution for each participant corresponds to a low-resolution mesh of approximately 32,000 vertices per hemisphere, mirroring the standard space for fMRI analysis in the Human Connectome Project (Glasser et al., 2013a).

### 2.7 CIFTI fMRI data generation

Preprocessed OC fMRI time series underwent surface mapping via Ciftify’s *ciftify_subject_fmri* pipeline, which uses a ribbon-constrained sampling procedure through the Connectome Workbench command line utilities (Glasser et al., 2013b). The resulting surface-mapped time series were spatially smoothed using a geodesic Gaussian kernel of σ = 4mm and excluded non-gray matter tissue.

### 2.8 Motion group classification

As expected, there was higher motion overall and larger inter-individual variation in the number of censored volumes in children relative to adults. To help distinguish effects of age from confounding effects of motion, we created groups of high- and low-motion children and applied the same threshold to adults. To generate a cutoff point, we examined the distribution of post-censored data for adults and children. We calculated and plotted probability density functions for each age group using their respective means for the full range of post-censored volumes (Supplemental Figure 1) and found the intersection of these curves occurred at 3863 volumes. Participants below this threshold were classified as the low-motion group and those exceeding this threshold were classified in the high-motion group. This resulted in 22 low-motion adults (LMA) (11 M, 11 F; aged 33.75 – 47.13 years; mean FD: 0.048 – 0.163), 10 low-motion children (LMC) (4 M, 6 F; aged 6.56 – 8.92 years; mean FD: 0.050 – 0.270), and 14 high-motion children (HMC) (8 M, 6 F; aged 7.03 – 8.90 years; mean FD: 0.122 – 0.939). An additional two high-motion adults (HMA) were excluded from analyses that used these motion groups but were retained for analyses where head motion was treated as a continuous variable. Sex, age, and head motion were compared across motion groups. A chi-squared test was used to examine sex differences. One-way ANOVAs were used to compare motion groups on age, censored volumes, and scan duration required to reach 30-minutes of post-censored data, along with post-hoc Tukey’s HSD tests to determine specific motion group differences.

### 2.9 Identification of networks in individuals using template matching

We used template-matching (Seitzman et al., 2019) to delineate individual functional networks for each participant and derive ROIs that had consistent network membership across the sample. This ensured that features assigned to a particular network are aligned with functional boundaries across age groups. Each cortical vertex was assigned to one of 14 functional networks based on the approach described in Gordon et al. (2017). We used a data-driven group network map, using vertex time series data derived from the Midnight Scan Club sample in previous work from our group (Rai et al., 2024), as our template, which was first binarized to the network labels from the WashU 120 map (Seitzman et al., 2019) (Supplemental Figure 2). For each participant, 60 minutes of post-censored data across all viewing conditions and sessions were used for template matching. Once individual network maps were created, we then computed a consensus adult group map and a consensus child group map using a 66% threshold (Supplemental Figure 3A). Next, the overlap between the child and adult maps created a final consensus network map for our sample (Supplemental Figure 3B). We labelled brain networks based on the established nomenclature from existing literature (Dworetsky et al., 2021; Laumann et al., 2015; Power et al., 2011) as: default mode (DMN), frontoparietal (FP), visual, dorsal attention (DAN), ventral attention (VAN), salience (SAL), auditory (AUD), cinguloopercular (CON), somatomotor dorsal (SMd), somatomotor lateral (SMl), temporal pole (Tpole), medial temporal lobe (MTL), parietal medial (PMN), and parietal occipital (PON).

### 2.10 High probability regions of interest (ROIs)

Using the MNI coordinates of 153 high probability regions of interest (ROIs) (Dworetsky et al., 2021), we first created volume spheres via Ciftify’s *ciftify_vol_result* command line function. Next, we projected those volume spheres into 3 mm cortical surface spheres using Ciftify’s *ciftify_surface_rois* function. We manually assessed the overlap between all surface spheres and the final consensus network map, which resulted in a total of 143 ROIs after excluding ROIs that did not overlap with the final consensus network map in our sample (Supplemental Figure 3C). We further excluded the PMN since network-level analyses required at least two ROIs within each network to compute averages. All other networks consisted of at least 2 ROIs for averaging, resulting in a final set of 142 ROIs across 12 networks.

### 2.11 Whole-connectome FC reliability

To assess whole connectome reliability as a function of scan duration, we used two approaches with the full post-censored data available across all viewing conditions and sessions. First, we employed a “split-session” approach, where we concatenated each participant’s time series data in three ways: session 1 and 4 vs. session 2 and 3, session 1 and 2 vs. session 3 and 4, and session 1 and 3 vs. session 2 and 4. In this approach, temporal sequencing of post-censored volumes is maintained. We computed FC matrices using Pearson correlations, applied Fisher z-transformation, and calculated test-retest correlations (TRC) by correlating the flattened matrices from each split. We evaluated FC-TRC using scan lengths from 5 to 80 minutes, in 5-minute increments, averaged across split combinations. We used a linear mixed-effects model to assess how age group, head motion, and their interaction affected the scan time required to reach a reliability threshold of FC-TRC ≥ 0.8, with family as a random effect. Following this, we performed a *post-hoc* analysis of the interaction effect by calculating slopes for the relationship between head motion and time-by-reliability for each age group. T-tests were then conducted to assess the significance of these slopes within each age group. Effect size was calculated using Cohen’s D for each time increment to show the magnitude of motion group differences in FC reliability. The incremental benefit was computed as the difference in FC-TRC between successive 5-minute scan lengths, for each motion group.

Second, we used an “iterative” approach, similar to Gordon et al. (2017) and Laumann et al. (2015) by correlating subsets of time series data. For each participant, we censored and concatenated data from all available sessions and randomly selected 1-minute chunks to make a 60 minute “true” reference subset of data and kept the remaining data as the “residual” subset. FC matrices were computed using Pearson correlations for each subset which were Fisher z-transformed, and FC-TRC was calculated by correlating the “true” data FC matrices with the “residual” data from 5 to 80 minutes in 5-minute intervals repeated across 1000 iterations and averaged. Here, data were randomly sampled from across the residual time series and therefore did not maintain temporal ordering of volumes. We again used a linear mixed-effect model to examine relationships between scan duration, head motion, and age group at a reliability threshold of FC-TRC ≥ 0.8, with family as a random effect. For comparison to the “split-session” approach, we also calculated effect sizes and incremental benefit of scan lengths as above.

### 2.12 Cortical surface FC reliability

We constructed parcellated FC connectomes across all viewing conditions and sessions to visualize test-retest correlations and ICC across the cortical surface for each motion group. Using the 1000-parcel Schaefer atlas (Schaefer et al., 2018) and Connectome Workbench’s *cifti-parcellate* command line utility, we parcellated each participant’s time series data across all sessions and viewing conditions. We matched each time series to include the participant with the lowest amount of usable data at 24 minutes of split-half data or 48 minutes of total data (i.e., 24 minutes from sessions 1 and 4 combined and 24 minutes from sessions 2 and 3 combined). Following this, we computed Pearson correlations to produce a 1000 x 1000 FC connectome for each participant within each motion group. We calculated intraclass correlation coefficient (ICC (2,1)) (Shrout & Fleiss, 1979) for functional connections of each parcel with all other parcels across participants separately for each motion group (LMA, LMC, HMC). We then averaged these ICC values across all connections for each parcel to visualize surface reliability maps for each motion group. To assess region-specific differences, we compared FC-TRC and ICC values between motion groups by subtracting the high- and low-motion children’s maps from the adults’ map.

### 2.13 Edge-wise FC reliability

To compare reliability across motion groups and time points, we calculated ICC matrices across all viewing conditions and sessions using varying split-scan durations based on available data for each motion group. For low-motion adults and low-motion children, we computed edge-wise ICC values at 5, 10, 24, and 54 split-half minutes (equivalent to 10, 20, 48, and 108 total minutes of data), and for the high-motion children we computed values up to 24 split-half minutes (48 total minutes of data). Edge-wise ICC values were computed, belonging to one of 12 networks, for each motion group. Next, we asked whether averaging edges within networks before computing ICC values would yield higher reliability compared to averaging post ICC computation within the same network regions. Following established conventions (Cicchetti & Sparrow, 1981), we categorized ICC values into four ranges: poor (0<ICC≤0.4), fair (0.4<ICC≤0.59), good (0.6≤ICC≤0.74), and excellent (ICC≥0.75).

### 2.14 Network-wise FC reliability in relation to scan duration

We considered time-by-reliability relationships in specific networks to assess whether adding more time always translated to more reliable data. We visualized specific networks with varying levels of reliability based on our edge-wise ICC findings from above. High reliability networks were DMN and FP, whereas lower reliability networks were AUD and SMd. For each network we computed ICC matrices at 5, 10, 24, and 54 minutes of split-half data for the LMA and LMC groups and only 5, 10, 24 minutes of split-half data for the HMC group due to data availability. We extracted each network’s ICC matrices and plotted the mean ICC value for each motion group at each time point.

### 2.15 Viewing condition analyses

To determine the variability in data retention between viewing conditions and motion groups, retention percentages were calculated for each condition by dividing the number of volumes retained by the total number of volumes collected (1640 volumes for each condition across all sessions). Repeated measures ANOVAs were conducted across motion groups, viewing conditions, and their interaction to assess differences in data retention. We performed *post-hoc* Tukey’s HSD tests for comparisons of data retention between motion groups and viewing conditions. A *post-hoc* pairwise t-test was also computed to assess significant differences within the motion group by condition interaction term, with Bonferroni correction applied to correct for multiple comparisons. Using the drowsiness and attention metrics, we performed a repeated measures ANOVA to examine the effects of viewing condition, motion group, and their interaction on drowsiness and attention scores. *Post-hoc* Tukey’s HSD tests were conducted to identify specific motion group differences in viewing conditions. We further conducted Levene’s tests to examine the variances in ratings across different conditions for both drowsiness and attention measures.

To compare FC-TRC across the different viewing conditions, we used the split-session approach. First, for each participant, we concatenated data from each condition separately in 1-minute increments and computed FC matrices across session 1 and 4 vs. session 2 and 3, sessions 1 and 2 vs. session 3 and 4, and session 1 and 3 vs. session 2 and 4. Next, we Fisher z-transformed connectomes and calculated FC-TRC as above for scan lengths from 1 up to 22 minutes. Using a two-way mixed ANOVA, we compared FC reliability across viewing conditions at 5-minutes. The ANOVA included age group (adult/child) as a between-subjects factor and viewing condition as a within-subjects factor, along with an interaction between age group and viewing condition. *Post-hoc* analysis included a paired t-test between viewing conditions, with Bonferroni-correction applied for multiple comparisons.

## 3 Results

### 3.1 Variability between motion groups

Sex did not vary significantly across the three motion groups (p > 0.05, chi^2^ = 1.80) and age did not differ between the low-motion and high-motion child groups (p > 0.05). Different amounts of scan time were required between motion groups to achieve 30 minutes of usable (post-censored) data. We required an average of 32.93 minutes (range: 30.49 – 37.18 mins) for the LMA group, 34.68 minutes (range: 30.67 – 37.28 mins) for the LMC group, and 50.85 minutes (range: 39.66 – 67.49 mins) for the HMC group (Figure 1). The high-motion adult group consisted of 2 participants, with an average of 61.47 minutes of data required to reach 30 minutes of usable data, which were not included in further group analyses. A one-way ANOVA revealed significant differences in data required between the three motion groups (F = 58.635, p < 0.001, ηp² = 0.732). Post-hoc Tukey’s HSD tests showed that the HMC group required significantly more data compared to both the LMA and LMC groups (p < 0.001).

**Figure 1:**
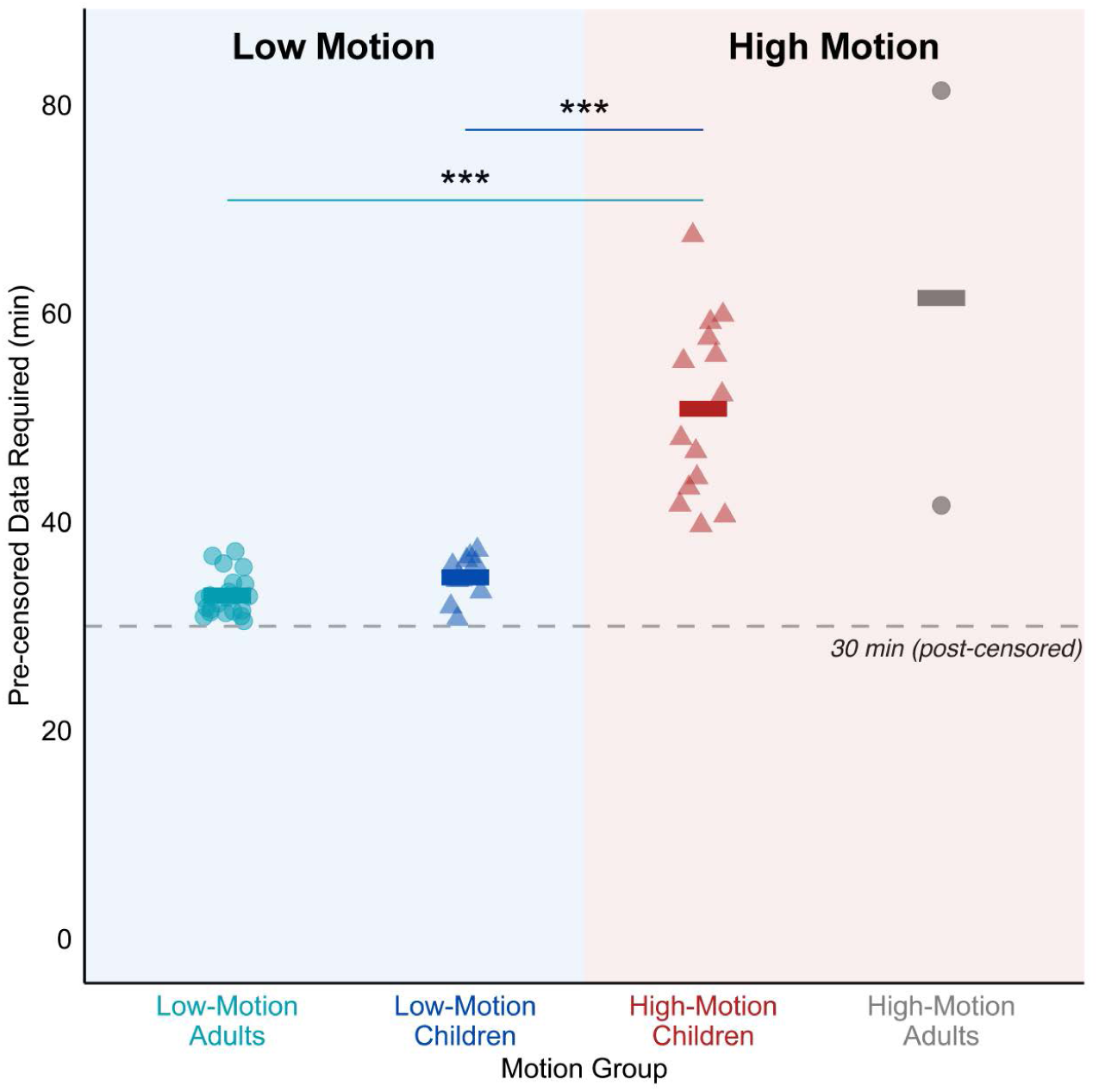
Pre-censored data required for 30 minutes of useable data using all viewing conditions across motion groups. The plot shows the total amount of data required to obtain 30 minutes of post-censored data (FD > 0.15mm) for each motion group. Low-motion adults (LMA, light blue circles) and low-motion children (LMC, dark blue triangles) required significantly less data compared to high-motion children (HMC, red triangles; both p < 0.001). High-motion adults (HMA, grey circles) were excluded from further motion group analyses. Each point represents a participant, and horizontal bars represent motion group means. The dashed line indicates the target 30-minute threshold of post-censored data.

As expected, based on our motion group definition, we found significant differences in censored volumes across the three motion groups (F = 84.701, p < 0.001, ηp² = 0.798) visualized in Supplemental Figure 4. *Post-hoc* tests revealed differences in censored volumes occurred between both the HMC and LMA groups (p-adjusted < 0.001) and between the HMC and LMC groups (p-adjusted < 0.001), but not between the LMA and LMC groups (p-adjusted = 0.223).

### 3.2 FC reliability with increasing scan duration

Across both approaches to calculating time-by-reliability curves, FC reliability asymptotically increased with scan length for all participants (Figure 2AB). Using the split-session approach (Figure 2A), we found that head motion (indicated by the color gradient) had a significant positive association with time to reach FC-TRC ≥ 0.8 (Standardized β = 0.531, SE = 0.158, z = 3.360, p = 0.001). Age group had an additional positive association with time to reach FC-TRC ≥ 0.8, with adults having shorter time to reach this reliability level (Standardized β = -0.993, SE = 0.236, z = -4.212, p < 0.001). We also found a significant interaction between head motion and age group (Standardized β = -0.603, SE = 0.241, z = -2.503, p = 0.012). Post-hoc analysis showed a significant positive effect of head motion on the time to reach FC-TRC ≥ 0.8 for children (SE = 0.158, t = 3.360, p = 0.002) and no significant effect for adults (SE = 0.183, t = -0.394, p = 0.696). Supplemental Figure 5A shows averaged reliability for each motion group. The amount of pre-censored data required to reach FC-TRC ≥ 0.8 was 15.82 minutes for the LMA group (range = 5.94 to 27.74 min), 23.74 minutes for the LMC group (range = 10.63 to 36.04 min), and 47.10 minutes for the HMC group (range = 22.13 to 78.74 min).

**Figure 2:**
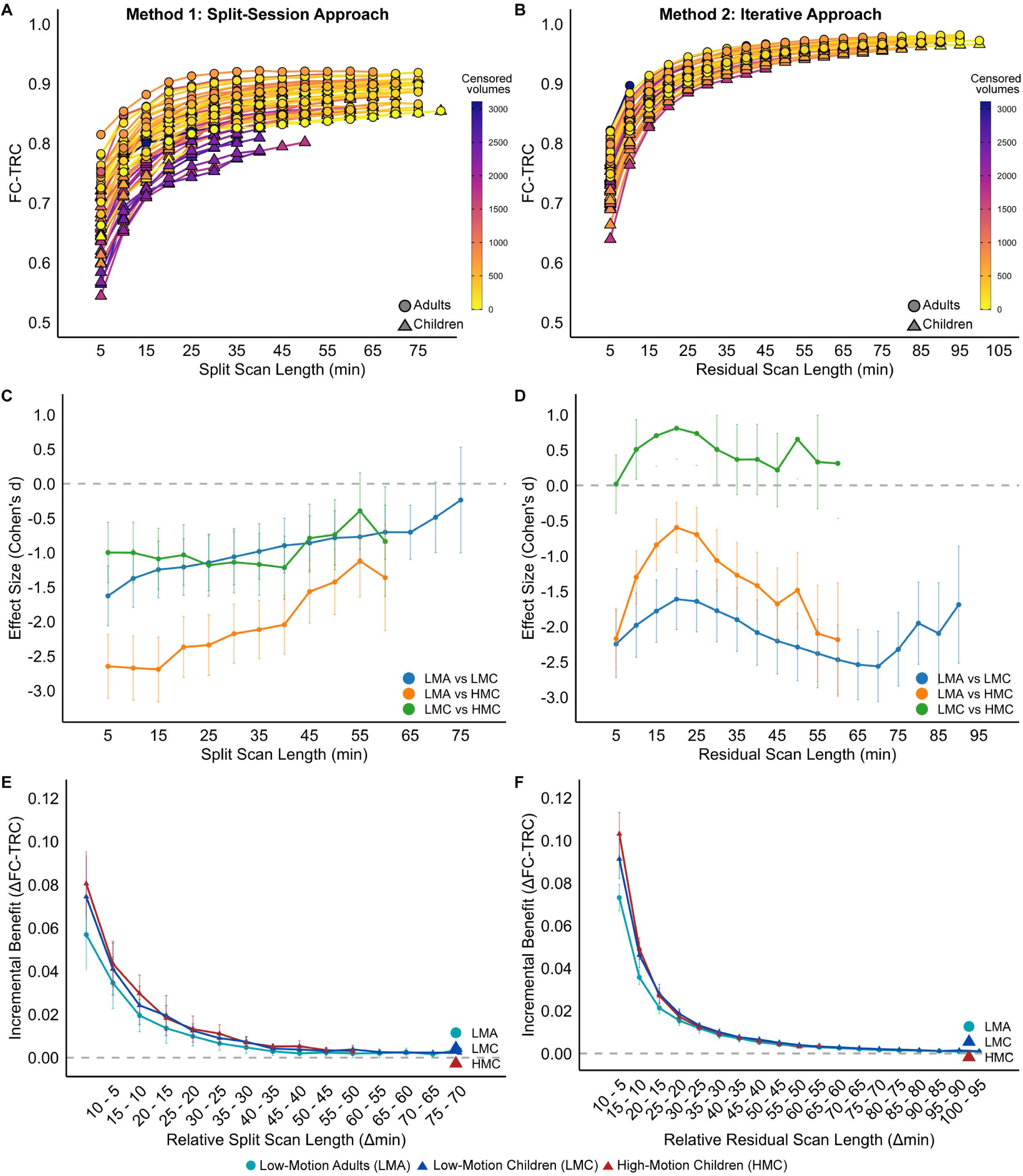
FC reliability asymptotically increase with scan duration using two methodological approaches under passive viewing conditions. Both the split session (A) and iterative (B) approaches show improved test-retest correlations (FC-TRC) as scan time increases, with color gradients indicating censored volumes (FD > 0.15mm). Both approaches highlight variations between adults and children to achieve an FC-TRC value ≥ 0.8. C) Effect sizes (Cohen’s d) for the split-session approach show the largest gap between low-motion adults (LMA) and high-motion children (HMC), with group differences diminishing at longer scan durations. D) Effect sizes for the iterative approach show no distinctions between the high-motion children (HMC) and low-motion children (LMC), and a less clear relationship with scan duration. E,F) The incremental benefits of FC-TRC relative to scan length show the most substantial advantage occurs between the 5 to 10-minute range, with larger gains for children relative to adults.

Using the iterative approach to calculate FC reliability (Figure 2B), we found a significant effect of age group on time to reach FC-TRC ≥ 0.8 (Standardized β = -1.645, SE = 0.235, z = -6.988, p < 0.001), but no significant effect of head motion (Standardized β = - 0.163, SE = 0.152, z = -1.077, p = 0.281) or an interaction between age group and head motion (Standardized β = -0.191, SE = 0.238, z = -0.802, p = 0.423). Averaged FC-TRC for all three motion groups are shown in Supplemental Figure 5B.

We next asked whether the reliability gap between adults and lower-and higher-head motion children closes when sufficient data are collected. Figure 2CD shows the effect size for differences in reliability between motion groups for different scan durations. Using the split-session approach (Figure 2C), we note that up to the largest amounts of split-half data available, the LMA group had higher reliability than either child group, but that more data reduced the gap between motion groups, with all pairwise effects trending towards zero. Using the iterative approach (Figure 2D), the data show more stable reliability gaps between motion groups, which do not converge with increasing amounts of data, and reliability curves distinguishing adults from children, independent of motion group.

Given the asymptotic shape of reliability curves, with diminishing reliability gains as scan length increased, we plotted the incremental reliability gains in Figure 2EF. For both methods, the benefits of collecting longer time series are higher for children than adults at the lowest amounts of data (e.g. going from 5 - 15 minutes). The curves converge and become essentially flat at 40 minutes for the split-session approach and 50 minutes for the iterative approach.

### 3.3 FC-reliability across the cortical surface

In Figure 3, with equally matched and post-censored data, the LMA group showed uniformly high reliability across the cortex, except within high-dropout ventromedial prefrontal and temporal pole regions. Greater heterogeneity across the brain was observed in both the LMC and HMC groups with the lowest FC-TRC values overall in the HMC group. Differences in reliability were small and relatively uniform across the cortex for the LMA and LMC groups. Differences between the LMA group compared to the HMC group were larger, particularly in regions of the ventral prefrontal cortex and temporal lobe, areas typically associated with the frontoparietal, default mode, and limbic networks.

**Figure 3:**
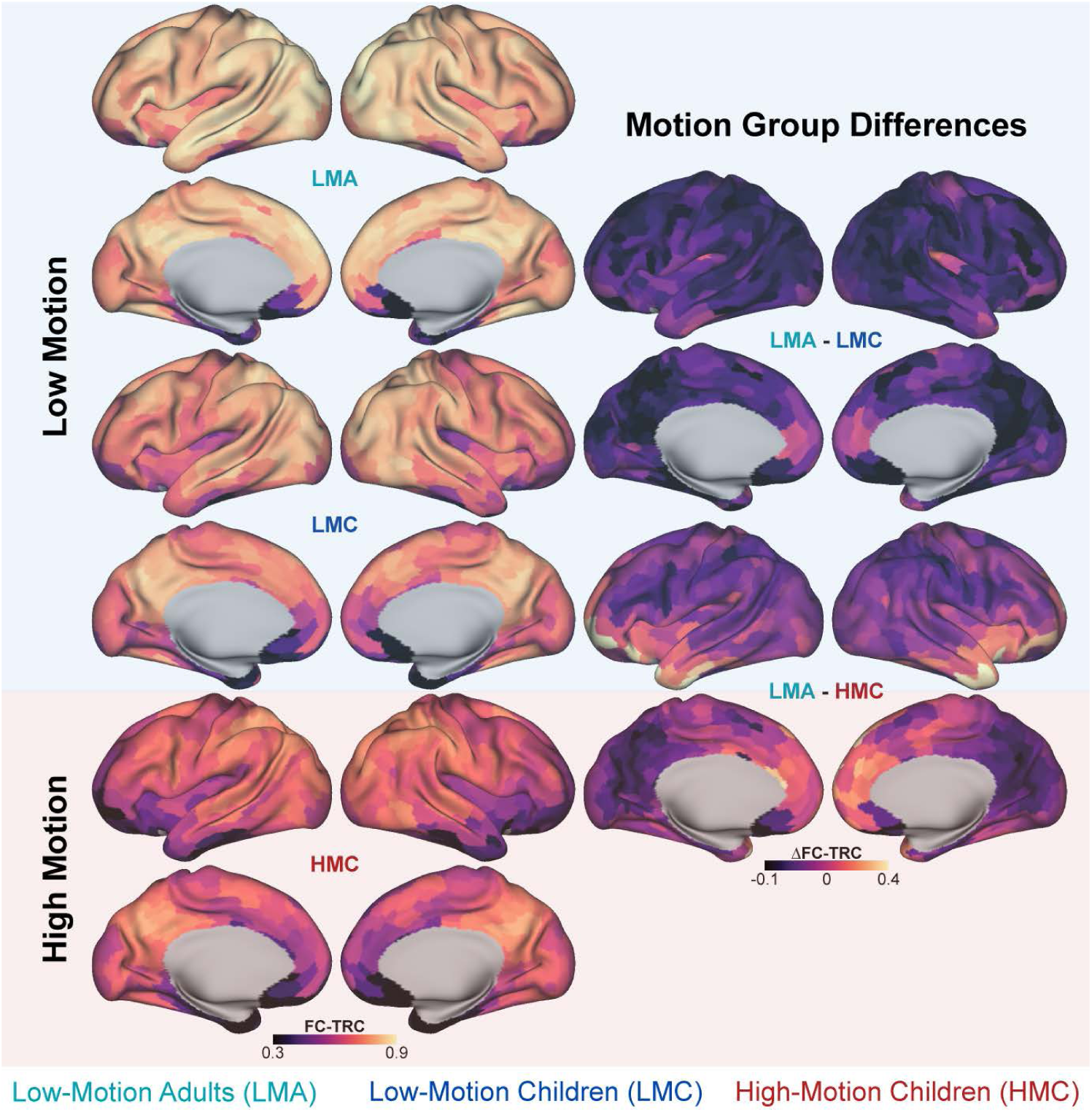
Regional differences in FC reliability across motion groups under passive viewing conditions. Cortical surface maps illustrate regional variations in test-retest correlations (FC-TRC) across motion groups using 24 minutes of split-half data. The left column shows FC-TRC maps for each motion group, with low-motion adults (LMA) demonstrating higher overall FC-TRC than both child groups, particularly in the precuneus, superior frontal lobe, and inferior parietal lobe. The right column features between-group difference maps, highlighting disparities between the low-motion adults (LMA) and high-motion children (HMC) within the frontoparietal, default mode, and limbic networks. Brighter colors represent regions with the most pronounced differences in FC-TRC between groups. The Schaefer 1000 parcel 17-network atlas was used to parcellate brain regions, and the split-session approach was used to compute FC-TRC.

Specifically, using the Schaefer 17-network atlas, regions exceeding a difference greater than 0.2 units between the LMA and HMC groups were parcels from the Control B (also known as frontoparietal network), Limbic A and B, Default A and B, and Salience/Ventral Attention A networks. While these observations are broadly consistent using ICC reliability measures in Supplemental Figure 6, the ICC analysis revealed more pronounced differences between motion groups.

### 3.4 Edge-wise FC-reliability by scan duration

Next, we assessed whether some functional network edges are more reliable with smaller amounts of data and whether acquiring more data improves reliability uniformly or focally (Figure 4). With 5 minutes of post-censored split-half data, specific edges within the DMN, FP and visual networks exhibited reliability values in the “good” to “excellent” ICC range across all three motion groups. With 24 minutes of split-half data, inter-network edges remained in the “poor” and “fair” ranges across all motion groups with fewer “good” to “excellent” reliability edges overall (both within and between) in the HMC group. With 54 minutes of split-half data, available only for the LMA and LMC groups, we observed global improvements in reliability both within and between most edges across both motion groups. Comparing ICC values for average network edges (Figure 5A) with the average edge-wise ICC values for the same networks (Figure 5B), we found that averaged network values generally had higher ICC values, supporting the use of network-averaged features from a reliability perspective.

**Figure 4:**
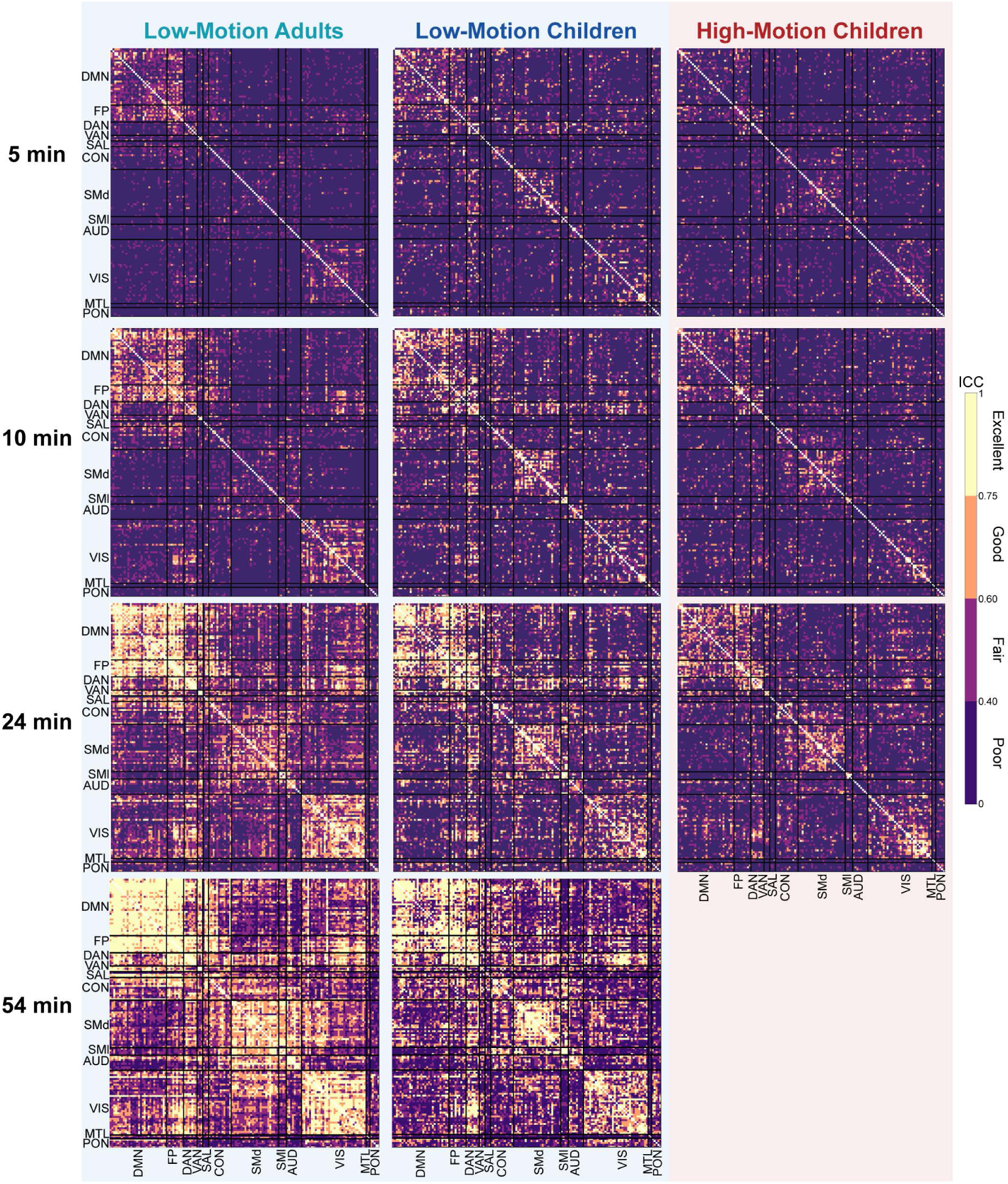
Edge-wise Intraclass Correlation Coefficient (ICC) reliability across motion groups and scan durations under passive viewing conditions. Matrices show ICC reliability for edge connections at 5, 10, 24, and 54 minutes of split-half data. Some edges show good-to-excellent reliability at 5 minutes of split-half data, especially within the default mode network, frontoparietal, and visual networks. As split-half scan duration increases, all groups demonstrate increases in edge-wise ICC reliability, with the most pronounced improvements noted in the low-motion adults (LMA) (left column). High-motion children (right column) demonstrate consistently lower ICC reliability across all scan durations. ICC values are color-coded as follows: poor (0<ICC≤0.4, dark purple), fair (0.4<ICC≤0.59, violet), good (0.6≤ICC≤0.74, orange), and excellent (ICC≥0.75, yellow).

**Figure 5:**
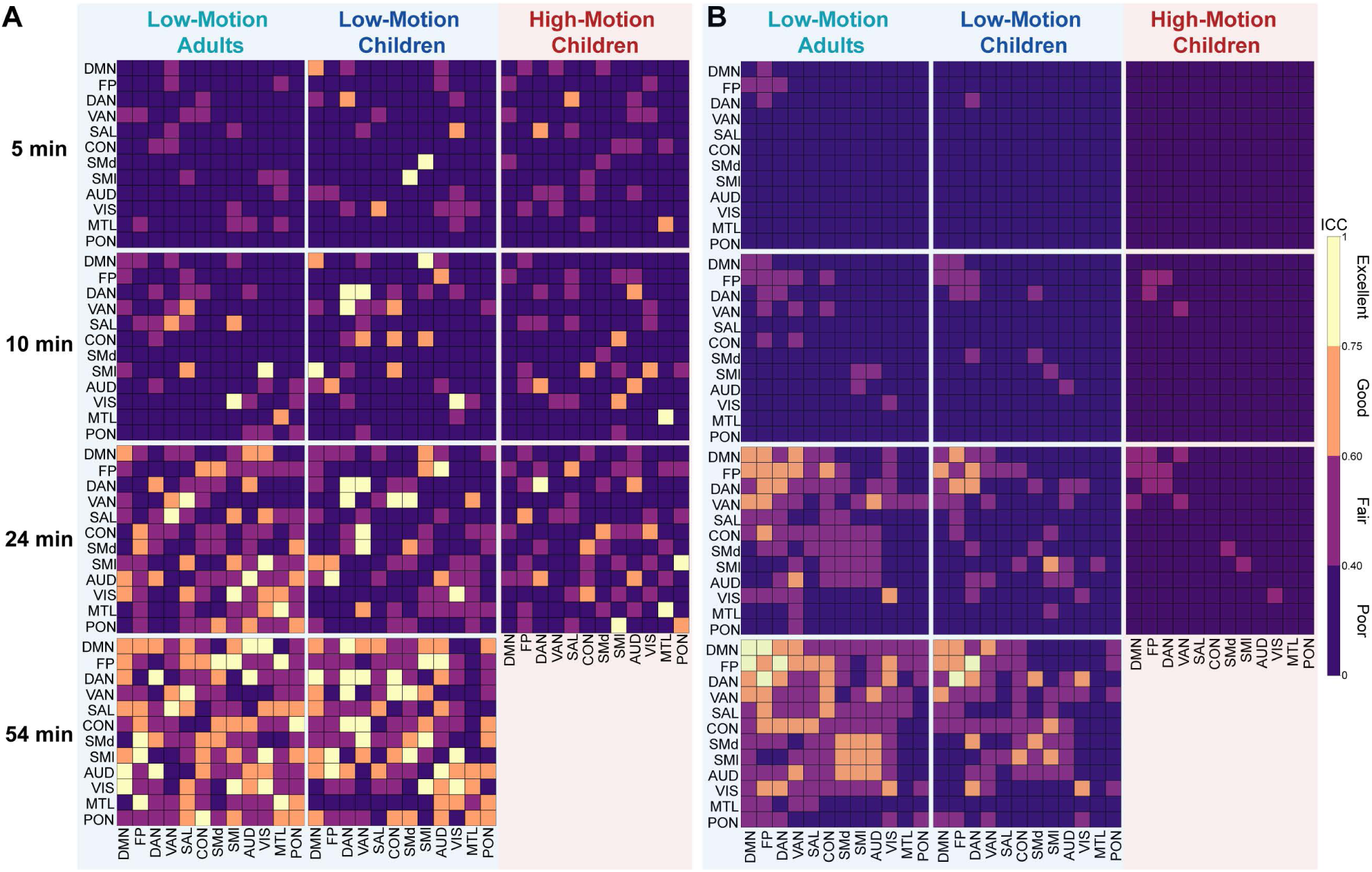
Network-wise Intraclass Correlation Coefficient (ICC) reliability comparing pre- and post-averaging approaches under passive viewing conditions. Matrices show ICC reliability across motion groups at 5, 10, 24, and 54 minutes of split-half data. (A) Network-wise values averaged prior to ICC computation generally resulted in relatively stronger ICC strengths relative to values averaged in the same networks post ICC computation in panel (B). ICC values are color-coded as follows: poor (0<ICC≤0.4, dark purple), fair (0.4<ICC≤0.59, violet), good (0.6≤ICC≤0.74, orange), and excellent (ICC≥0.75, yellow).

### 3.5 Network-wise FC-reliability by scan duration

We next considered specific functional networks with average high and low reliability features, to ask whether increasing scan time uniformly increases reliability (Figure 6). Specifically, we selected DMN and FP as the higher reliability networks and AUD and SMd for our lower reliability networks. Overall, ICC ranges improved with increasing scan duration, but for some networks the gap in reliability between the lowest and highest motion participants remained relatively stable (DMN, AUD), while for others the gap was smaller and less consistent across scan durations (FP, SMd). This suggests that while increasing scan length is generally beneficial for edge-wise reliability, it does not always overcome residual effects of signal drop-out, head motion and censoring on the BOLD time series.

**Figure 6:**
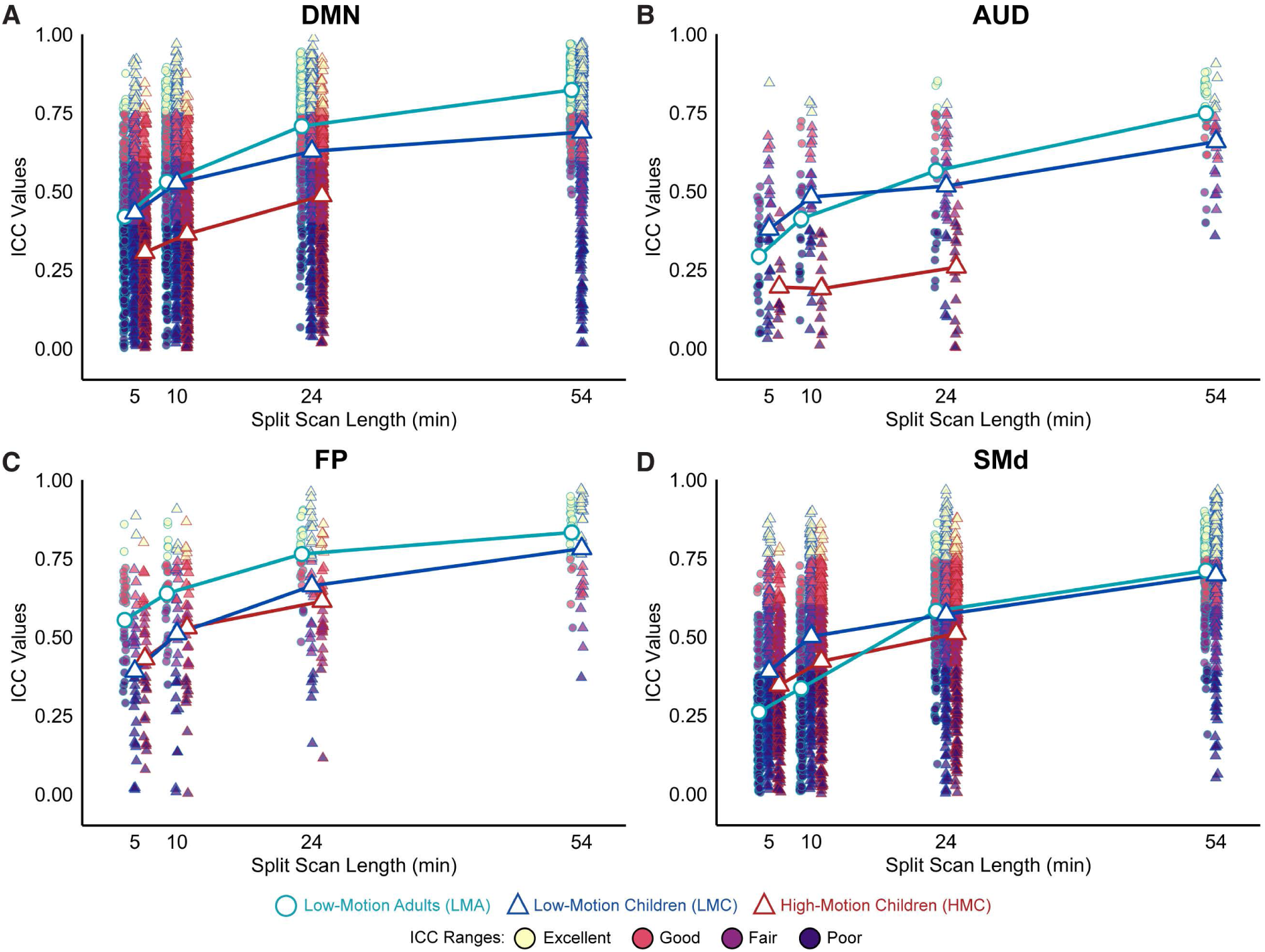
Intraclass Correlation Coefficient (ICC) reliability in specifically selected high reliability networks (DMN, FP) and low reliability networks (AUD, SMd) during passive viewing. A,B) The difference between groups in ICC reliability remain large between low motion and high motion groups for the DMN and AUD networks. In contrast, in (CD) the difference observed in the FP and SMd networks is less widespread. Low-motion adults (LMA, light blue circles) consistently exhibited higher ICC values compared to both low-motion children (LMC, dark blue triangles) and high-motion children (HMC, red triangles). ICC values are color-coded as follows: poor (0<ICC≤0.4, dark purple), fair (0.4<ICC≤0.59, violet), good (0.6≤ICC≤0.74, orange), and excellent (ICC≥0.75, yellow).

### 3.6 Participant engagement and data retention across viewing conditions

Participant drowsiness was significantly associated with viewing condition (F = 6.95, p = 0.001, ηp² = 0.139) (Figure 7A), but there was no significant effect of motion group (F = 2.96, p = 0.055) or an interaction between motion group and condition (F = 0.48, p = 0.753). Post-hoc analyses found self-reported drowsiness scores were significantly higher in the low-demand condition compared to both narrative (p-adjusted = 0.003) and non-narrative conditions (p-adjusted= 0.007). Attention scores also significantly associated with viewing condition (F = 9.11, p < 0.001, ηp² = 0.175) (Figure 7B), but there was no significant effect of motion group (F = 1.78, p = 0.172) or motion group by viewing condition interaction (F = 0.59, p = 0.673). Post-hoc tests showed that attention scores were significantly lower in the low-demand condition compared to both narrative (p-adjusted = 0.001) and non-narrative (p-adjusted < 0.001) conditions. Levene’s tests revealed significant differences in variance for the drowsiness measure across conditions (F = 3.62, p = 0.029), with the low-demand condition showing the highest variance. Similarly, for the attention measure, there were significant differences in variances across viewing conditions (F = 11.74, p < 0.001), with the variance in attention scores for the low-demand condition again substantially higher than the other conditions.

**Figure 7:**
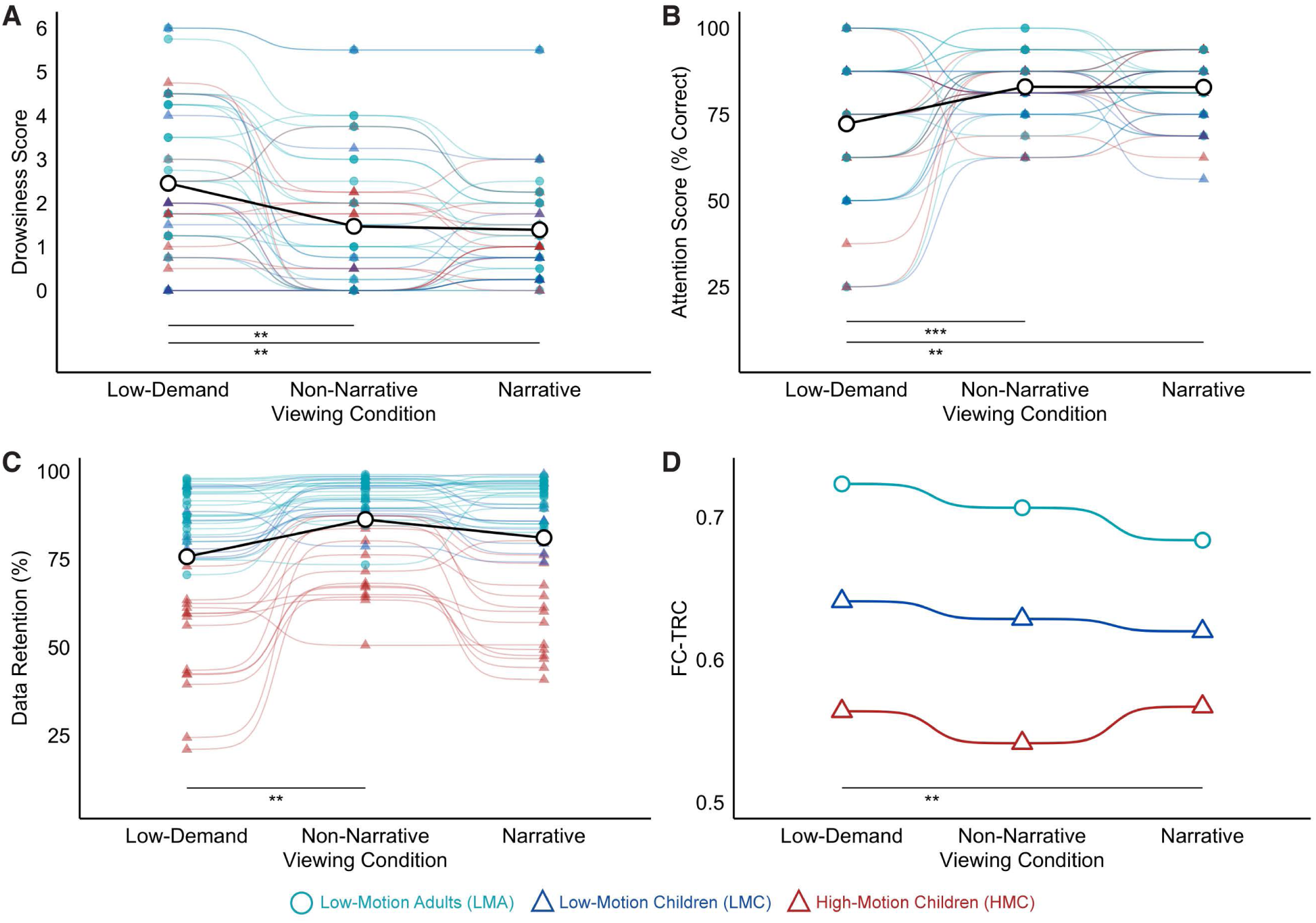
Trade-offs between participant engagement, data retention, and FC reliability across viewing conditions. A) Self-reported drowsiness levels were highest during the low-demand condition across all motion groups. B) Attention scores were significantly reduced during the low-demand condition compared to other conditions. C) Data retention percentages post-censoring showed significant differences between the low-demand and non-narrative conditions. D) FC-TRC, measured at 5 minutes of split-half data, was highest during the low-demand condition across motion groups and had significantly higher FC-TRC than the narrative condition. Low-motion adults (LMA, light blue circles) consistently showed better metrics compared to low-motion children (LMC, dark blue triangles) and high-motion children (HMC, red triangles). Significance: ** p < 0.05, *** p<0.001.

In terms of data retention, we observed significant differences between the motion groups (F = 84.701, p < 0.001, ηp² = 0.797), viewing conditions (F = 27.755, p < 0.001, ηp² = 0.392), and their interaction (F = 7.904, p < 0.001, ηp² = 0.269) in Figure 7C. *Post-hoc* analyses found that the HMC group had significantly lower data retention compared to both LMA (p-adjusted < 0.001) and LMC groups (p-adjusted < 0.001), with a difference of 30.8% and 26.3% respectively, while the LMA and LMC groups did not differ significantly (p-adjusted = 0.123). When considering viewing condition, *post-hoc* analyses found that the non-narrative condition had significantly higher data retention than the low-demand condition (p-adjusted = 0.008), with a difference of 10.6%. Retention differences between the other conditions were not statistically significant (p-adjusted > 0.05). When considering the motion group by condition interaction, we found that the HMC group had significantly lower retention compared to both LMA and LMC groups across all conditions (all p-adjusted < 0.001), with the largest differences found in the low-demand condition (HMC vs. LMA: 37.8%, p < 0.001; HMC vs. LMC: 32.8%, p-adjusted < 0.001) and smallest in the non-narrative condition (HMC vs. LMA: 20.2%, p < 0.001; HMC vs. LMC: 18.5%, p-adjusted < 0.001). We found no significant differences between LMA and LMC groups for any condition (all p-adjusted > 0.09). In sum, these results indicate that the choice of viewing condition impacts both variation in participant engagement and data retention.

### 3.7 FC reliability based on viewing condition

Finally, we considered the effect of viewing condition on averaged FC reliability across motion groups using only the split-session approach (Supplemental Figure 7) as there was insufficient post-censored data to use the iterative approach. We assessed significant reliability differences using 5 minutes of split-half data and again found significant main effects of age group (F = 42.056, p < 0.001, ηp² = 0.662). More notably, in Figure 7D, we found significant effects of viewing condition on FC reliability with 5 minutes of split-half data (F = 4.197, p = 0.018, ηp² = 0.089). Conducting pairwise comparisons between viewing conditions, we found the only significant difference occurred between the narrative and low-demand conditions (t = -2.742, p-adjusted = 0.026, d = -0.265), with the low-demand condition showing higher FC reliability than the narrative condition. The group x condition interaction term was not significant (p-adjusted > 0.05).

## 4 Discussion

While adult studies have provided data on the relationship between scan time and FC reliability (Braga & Buckner, 2017; Gordon et al., 2017; Laumann et al., 2015), there are no current guidelines specific to pediatric samples. This is an important gap given the complexities involved with imaging children, including higher rates of head motion, difficulty remaining still for long scan durations, and the need for child-friendly viewing conditions (Barkovich et al., 2019; Raschle et al., 2012). Using repeated sampling of adults and pre-adolescent children across varying movie and video content stimuli, the present study considered FC reliability as a function of scan duration, head motion and viewing condition. Overall, our findings suggest that although reliability increases with scan time, *both* head motion and younger age can have a dampening effect on FC reliability even for relatively long durations. Age and head motion effects on reliability are not spatially uniform and may impact anterior/ventral regions most strongly. We also found that the choice of viewing condition significantly impacts both data retention and FC reliability. Together, our work suggests special consideration is needed when adapting study designs from adults to children.

We considered two approaches for calculating test-retest correlations, which we refer to as split-session and iterative. The iterative approach, used in previous adult studies (Gordon et al., 2017; Laumann et al., 2015), allows a separation of reliability from the timing and temporal sequence of volumes used for connectome calculation. The split-session approach arguably has more ecological validity and can replicate scenarios where, for example, only 10 minutes of data are collected. These two approaches provide different and complementary information about the factors influencing reliability. While the iterative approach is not sensitive to head motion, as might be expected based on randomly sampled volumes over many iterations, the split-session approach captures motion effects and suggests that motion interruptions at different times in a pair of runs reduce test-retest correlations. Both approaches show age effects, though in the split-session approach the reliability gap between groups decreased with increasing scan length, whereas the gap remained relatively stable in the iterative approach. Together, these results suggest that: 1) connectomes calculated from continuous and motion-free data (e.g., 10 minutes) may be more reliable than from the same amount of post-censored data, and 2) relatively lower whole connectome test-retest correlations in children cannot be solely attributed to head motion.

Several studies have examined FC reliability in pediatric populations (Somandepalli et al., 2015; Thomason et al., 2011; Y. Wang et al., 2021), though direct comparisons of FC reliability between children and adults remain limited. Our results of FC reliability differences between adults and children align with previous work showing age-related differences in task fMRI reliability (Kennedy et al., 2022; Koolschijn et al., 2011). We speculate that our findings represent a meaningful developmental difference, rather than solely due to group differences in head motion, as we know that brain structure and function changes across adolescence (Blakemore, 2008; Casey et al., 2008; Mills et al., 2016). These differences may reflect age-related differences in functional network topography (Bottenhorn et al., 2023; Cui et al., 2020) further evidenced by ‘fuzzier’ network boundaries in pre-adolescents (Tooley et al., 2022), which could ultimately translate to lower connectome reliability.

By considering FC-TRC as a function of scan duration, our findings provide information about reliability for shorter duration scans, as well as for pediatric precision fMRI applications. On average, children required nearly twice the post-censored scan time (24.6 minutes) compared to adults (14.4 minutes) to reach a predefined level of whole-connectome reliability (FC-TRC ≥ 0.8), which was achieved with more than double the amount of data (36.5 minutes of pre-censored data required for children and 16.9 minutes for adults). These findings underline the importance of minimizing head motion by using engaging stimuli (e.g., movies) (Finn & Bandettini, 2021; Greene et al., 2018; Meer et al., 2020; Vanderwal et al., 2015) or weighted blankets and incentive systems (Horien et al., 2020). Moreover, real-time motion monitoring systems like FIRMM (Dosenbach et al., 2017) could be calibrated for children to require longer amounts of motion-free data, given that motion effects reduce FC reliability even post-censoring (Satterthwaite et al., 2012; Van Dijk et al., 2012).

In terms of implications for precision fMRI, our study speaks to the feasibility of pfMRI (Gordon et al., 2017; Gratton, Kraus, et al., 2020; Michon et al., 2022) in pre-adolescent children during naturalistic conditions. Across four sessions, our study retained all recruited participants and found no significant differences in sex distribution or age between the low- and high-motion child groups. Our results do however suggest that at relatively long scan durations, reliability gaps between adults and children, particularly high-motion children, persist and that while volume censoring techniques can mitigate motion-related bias, (Ciric et al., 2017; Power et al., 2014), there may remain increased variance in these data (Parkes et al., 2018). Moreover, spatial differences in reliability suggest that even with longer scan durations, it may be challenging to precisely delineate specific functional networks in high-motion children.

Exploring group differences in spatial patterns of FC-TRC using 24 minutes of post-censored split-half data showed that adults had relatively uniform and high reliability across the cortical surface while children, especially high-motion children, exhibited greater spatial variance and lower overall reliability. The most prominent differences between adults and high-motion children were observed in ventral prefrontal and temporal regions, frequently associated with limbic networks that are prone to dropout. Our findings suggest that results pertaining to these regions should be interpreted with greater caution due to their lower reliability relative to other regions, particularly when examining age-related differences or longitudinal changes. We also note that while we used a multiband, multi-echo fMRI protocol for this study – acquisitions that have shown to improve temporal resolution and reduce susceptibility artifacts (Lynch et al., 2020) – this did not completely mitigate low reliability in high dropout regions.

Practically, with 5 minutes of post-censored data, we generally found poor ICC reliability across all groups and functional networks, aligning with prior work that suggests short scans result in poor overall network reliability (Noble et al., 2019a). We observed global improvements in reliability from 5 to 24 minutes of split-half data across groups, consistent with studies evaluating the benefits of longer scan durations (Anderson et al., 2011; Birn et al., 2013; Noble, Scheinost, et al., 2017; Shehzad et al., 2009; Termenon et al., 2016; Tian et al., 2021). Expanding on our previous finding on the effects of motion on reliability, here, we again found ICC reliability differences between age and motion groups even at 24 minutes of split-half scan time. At 54 minutes of split-half scan duration, a small gap between low-motion adults and low-motion children remained. Averaging functional network values prior to ICC computation, rather than post ICC computation, generally boosted reliability and complements previous findings (J. Wang et al., 2017a) establishing superior ICC ranges for averaged network values compared to unit-wise values, though most remained in the poor to fair range with 10 minutes of post-censored split-half data.

Though FC reliability analyses presented here examined the consistency of multivariate functional connectivity patterns using test-retest Pearson’s correlations (Damoiseaux et al., 2006) and univariate connectivity patterns using ICC (Brandmaier et al., 2018; Chen et al., 2018), there are additional multivariate approaches that can be employed. Measures such as discriminability and image intraclass correlation coefficient (I2C2) demonstrate higher reliability than univariate ICC (Camp et al., 2024; Noble, Spann, et al., 2017) and these multivariate metrics have also shown improved FC reliability in visual and temporal brain regions using movies compared to rest (Shearer et al., 2024). Future work should examine developmental differences in FC reliability by incorporating these complementary multivariate measures.

Our analysis of viewing conditions revealed significant effects on both data retention and FC reliability. Overall, in adults, both the narrative and non-narrative conditions had comparable post-censored data retained, while in children, the more engaging non-narrative condition retained the most data. Across all participants, the low-demand condition had the highest drowsiness levels and lowest attention scores, and the highest variation across participants. The impact of viewing condition on FC reliability was significant, albeit relatively small, with higher reliability in the low-demand condition relative to narrative and non-narrative conditions. While previous research has demonstrated that movies typically achieve similar or greater FC reliability compared to rest (Meer et al., 2020; J. Wang et al., 2017b; Zhang et al., 2022), and comparable ICC reliability across movies, Inscapes, and flanker task (O’Connor et al., 2017), more work is needed to directly compare FC reliability between videos with varying levels of engagement (Vanderwal et al., 2019). Recent work (Tian et al., 2021) across different movies found that whole-brain connectome ICCs were marginally higher for independent Creative Commons style of movies compared to commercial Hollywood style of movies, with relatively subtle differences in ICC within specific functional networks, which may be due to differential activation patterns based on the task at hand (Rai et al., 2024). Our findings expand and support previous research showing that engaging naturalistic paradigms enhance compliance, increase homogeneity of engagement levels, and reduce motion in developmental samples (Greene et al., 2018; Vanderwal et al., 2015). However, our findings indicate trade-offs between participant compliance and FC reliability.

The choice of viewing condition for pediatric fMRI can be challenging, and the options we compared here are far from exhaustive. While the choice of stimuli geared towards pre-adolescent children may limit the generalizability of our findings, we hope that this comparison provides some useful metrics for consideration of trade-offs in stimulus choice. We note that while the low-demand condition used here was inspired by Inscapes (Vanderwal et al., 2015), it is not identical to Inscapes or other nature-based passive viewing conditions (Vanderwal et al., 2019). Overall, the field can benefit from comparing naturalistic fMRI paradigms with the goal of identifying stimuli that balance between compliance, engagement, and reducing head motion (Finn, 2021).

## 5 Limitations

Strengths of this study include leveraging a unique dataset with >2.5h data collected in pre-adolescent children and well-matched adult controls. Limitations include a narrow pre-adolescent age range which could limit the generalizability of our findings. We also note that sample, data acquisition (including viewing condition) and processing choices (such as multi-echo ICA and global signal regression) all impact reliability (Anderson et al., 2011; Bennett & Miller, 2013; Cho et al., 2021; Graff et al., 2022; Murphy & Fox, 2017; Noble et al., 2019b; Noble, Spann, et al., 2017; Parkes et al., 2018; Rai et al., 2024; Ramduny et al., 2024; Tozzi et al., 2020). Therefore, specific values presented here will not apply directly to different study designs. While an exhaustive comparison of scan parameters, sample characteristics, and processing and methodological choices are beyond the scope of this work, we feel that our findings can nonetheless provide useful guidelines for adapting study designs to pediatric samples and add to the chorus that reliability considerations are important for reproducible neuroimaging findings (Bennett & Miller, 2010; Lynch et al., 2020; Vanderwal et al., 2019; Xu et al., 2023).

## 6 Conclusions

Using a unique dense sampling dataset, this study provides several practical considerations for fMRI-FC in children under passive viewing conditions. First, we demonstrated that dense sampling designs are feasible in pre-adolescent children, however, we emphasize the need for more extensive data collection for pediatric samples to achieve FC reliability comparable to adults. Relatedly, we also observed that reliability improvements show diminishing returns after 40-50 minutes of post-censoring data and that head motion may remain a persistent confound, such that differences in reliability remained substantial between lower and higher motion participants even with 24 minutes of split-half data (48 minutes of total scan duration). In terms of spatial patterns of reliability, our work suggests that head motion has a particularly detrimental impact on high-dropout regions. And lastly, our investigation of viewing condition showed nuanced trade-offs between data retention, participant engagement, and FC reliability. Together, our findings provide insights towards more reliable fMRI-FC in children.

## 7 Data and Code availability

The fully preprocessed and surface projected PreciseKIDS dataset, along with Python version 3.12.2 and MATLAB version 9.11.0 (R2021b) (The MathWorks Inc., 2022) scripts used in this study are available at https://github.com/BrayNeuroimagingLab/BNL_open.

## 8 Author Contributions

Shefali Rai: Conceptualization, Data Curation, Methodology, Validation, Formal analysis, Investigation, Writing - original draft, review & editing, Visualization, Funding acquisition. Kate J. Godfrey: Validation, Investigation, Writing - review & editing. Kirk Graff: Investigation, Writing - review & editing. Ryann Tansey: Investigation, Writing - review & editing. Daria Merrikh: Resources, Investigation, Writing - review & editing. Shelly Yin: Resources, Investigation, Writing - review & editing. Matthew Feigelis: Methodology, Investigation, Writing - original draft, review & editing. Damion V. Demeter: Methodology, Investigation, Writing - original draft, review & editing. Tamara Vanderwal: Conceptualization, Writing - original draft, review & editing. Deanna J. Greene: Methodology, Investigation, Writing - original draft, review & editing. Signe Bray: Conceptualization, Methodology, Investigation, Supervision, Project administration, Funding acquisition, Writing - review & editing.

## 11 Acknowledgements

We would like to acknowledge the dedication of all the families that participated in our study, along with the support of the Diagnostic Imaging staff at the Alberta Children’s Hospital.

## 12 Supplementary Material

All supplemental materials are provided separately.

