## Supplementary material for "How much is ‘enough’? Considerations for functional connectivity reliability in pediatric naturalistic fMRI": 12 Supplementary Material

**Supplemental Materials**

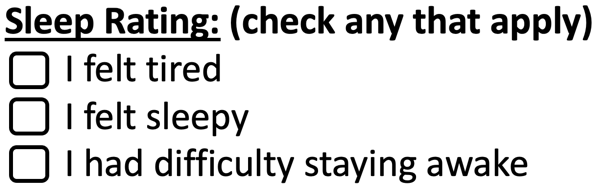
What was the name of the ancient lost city?

A) Papaya
B) Parapata
C) Pandora

How did Dora feel when Diego was leaving to move away?

A) Angry
B) Annoyed
A) Sad

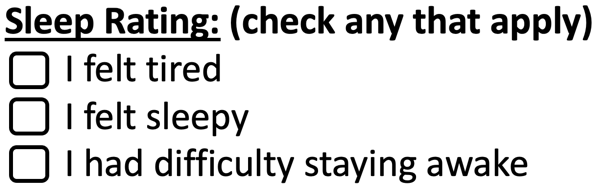
What kind of toy did the boy take out of the box?

A) Dinosaur
B) Legos
C) Fire Truck

How did the rainbow sand video make you feel?

A) Sad
B) Calm
C) Worried
D) Other: ____________

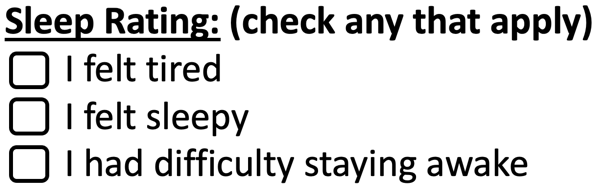
What season is the forest walking video showing?

A) Winter
B) Fall
C) Nighttime

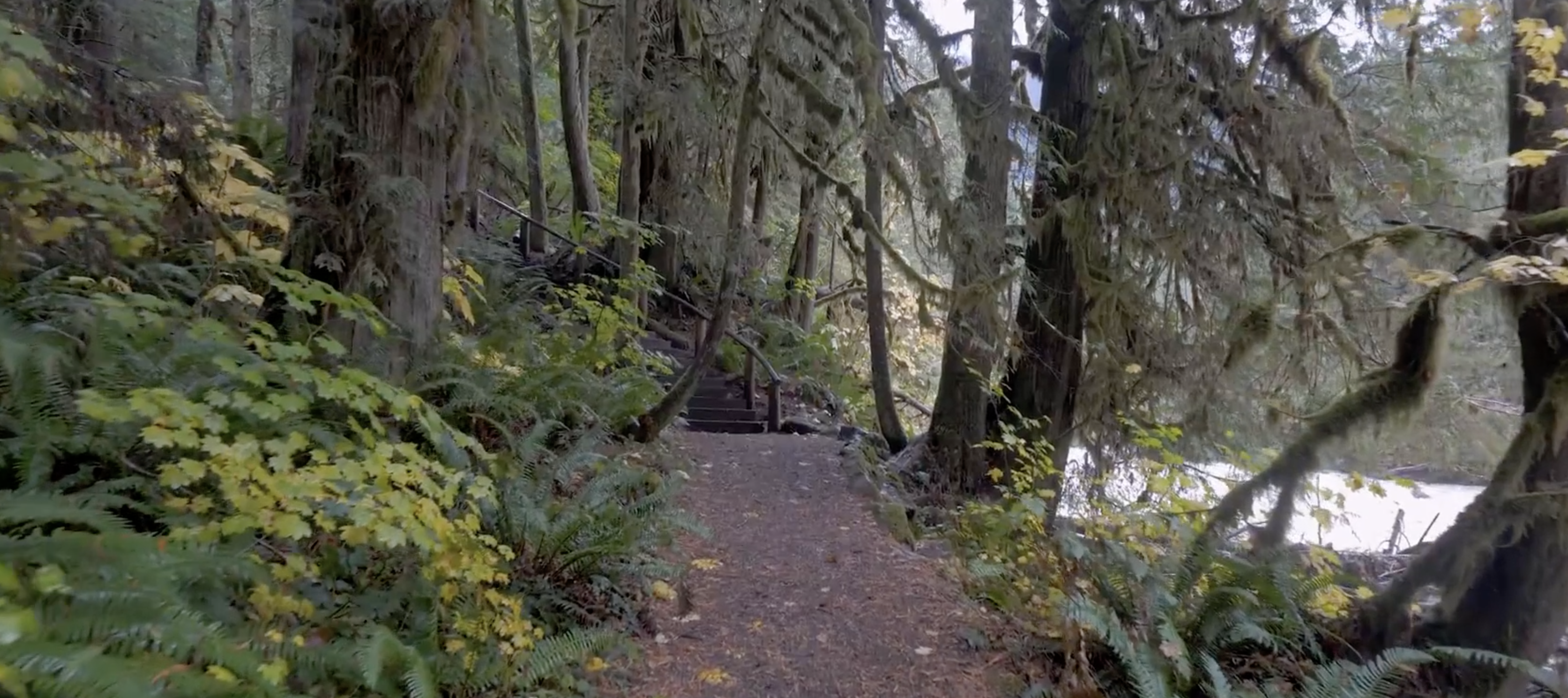

**Supplemental Table 1: Example post-MRI survey**

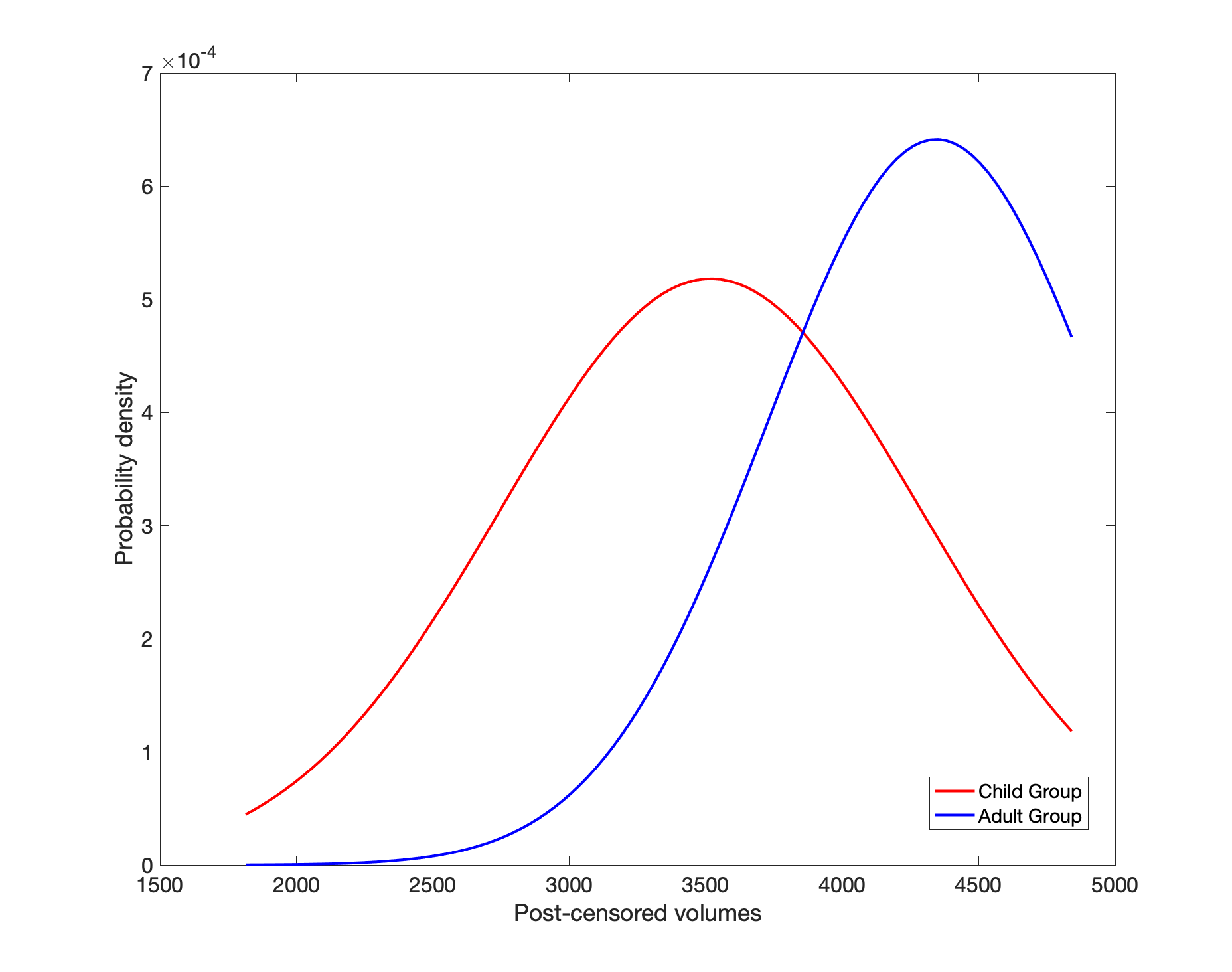

**Supplemental Figure 1: Probability density distribution of post-censored data acquired from adults and children during naturalistic fMRI.** Plotting the probability density across both groups, we observed an intersection of the curves at 3863 post-censored volumes. This point served as the threshold for classifying participants into the low-motion adults (LMA), low-motion children (LMC), and high-motion children (HMC) groups.

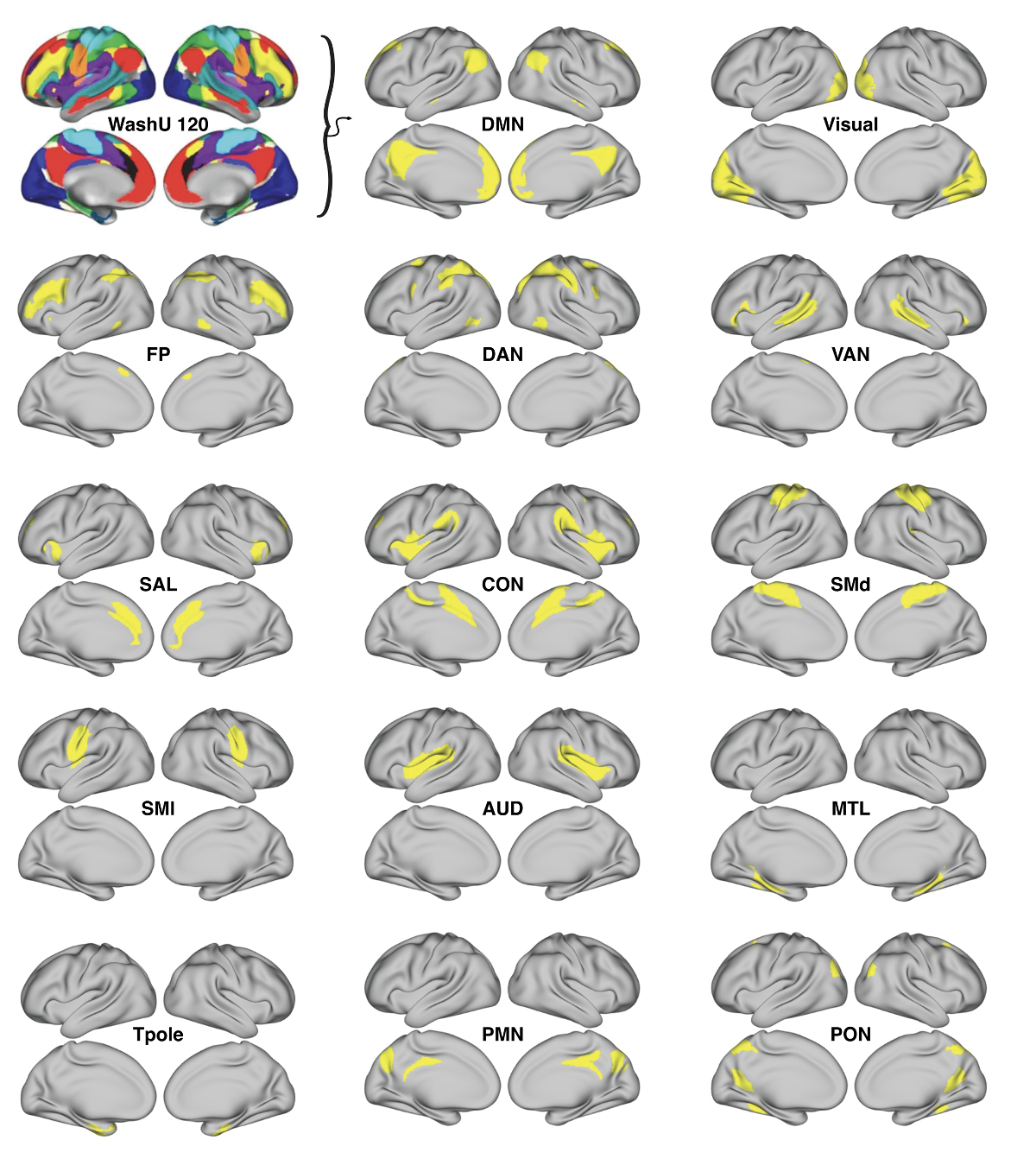

**Supplemental Figure 2: 14 network templates binarized to the WashU 120 template using vertex time series data from the Midnight Scan Club dataset, processed in a previous study from our group (Rai et al., 2024).**

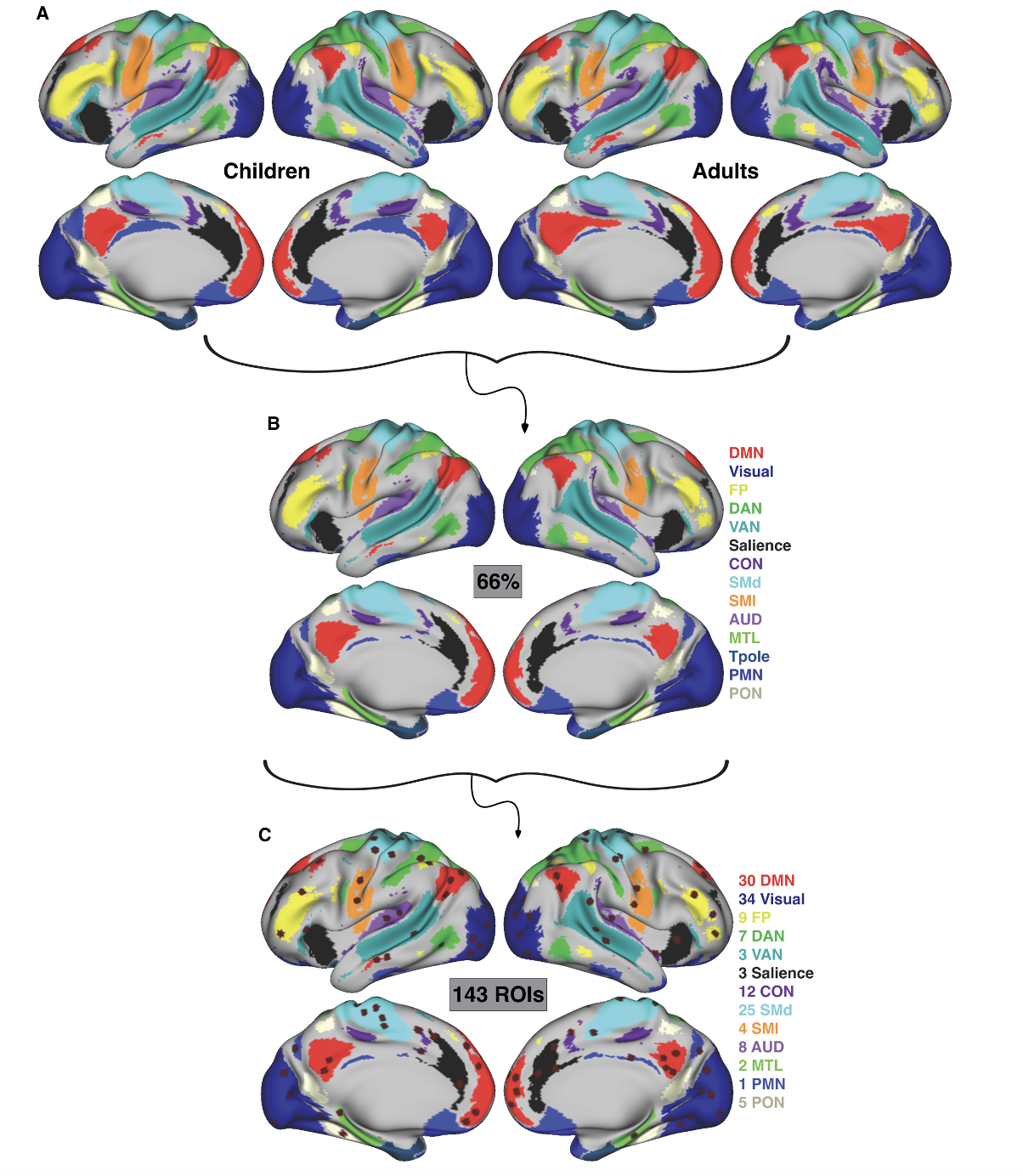

**Supplemental Figure 3: 143 consensus ROIs shown plotted on the combined adult and child group consensus atlas under passive viewing conditions.**

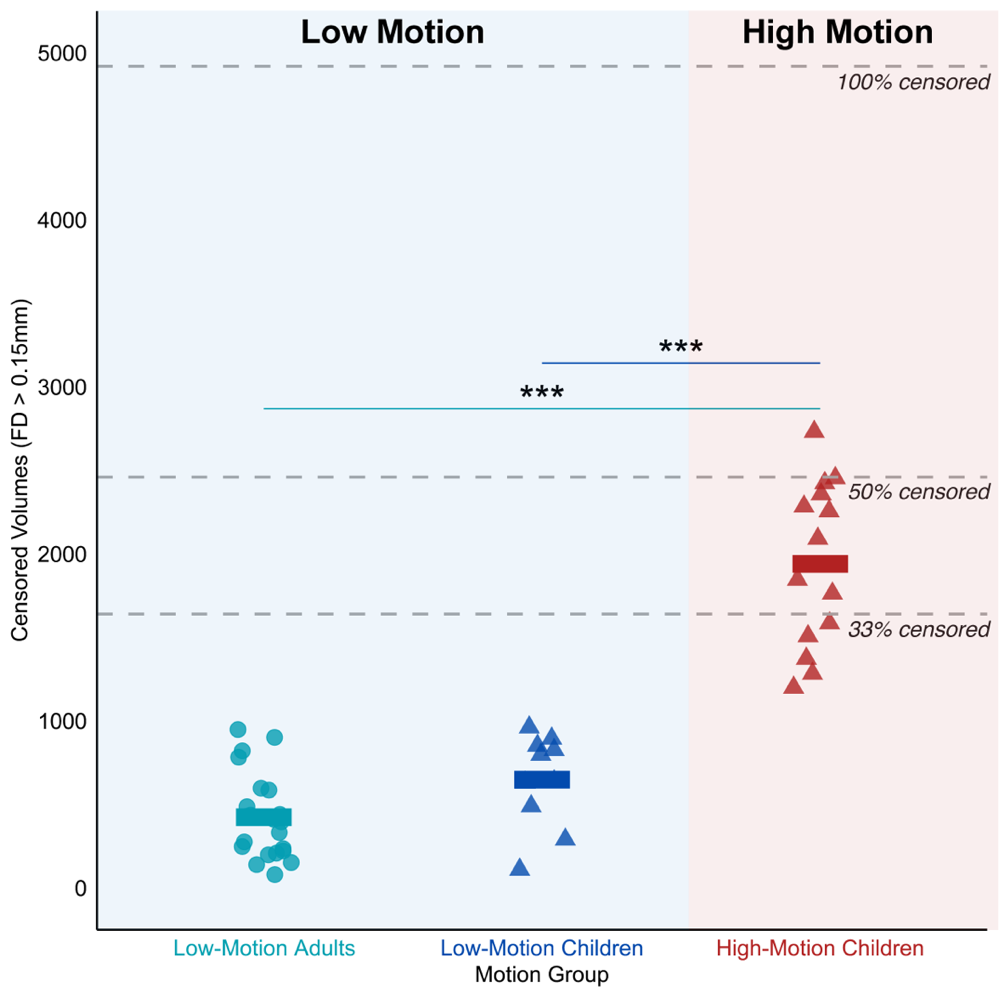

**Supplemental Figure 4: Amount of data censored across all passive viewing conditions for each motion group.** The number of censored volumes (FD>0.15 mm) were significantly different between the high-motion group (HMC, red triangles) and both the low-motion adults (LMA, light blue circles) and low-motion children (LMC, dark blue triangles). The low-motion adults did not significantly differ in censored volumes with the low-motion children (p-adjusted = 0.223). Dotted lines represent percent of volumes censored out of the 4920 total pre-censored volumes collected. Significance: *** p<0.001.

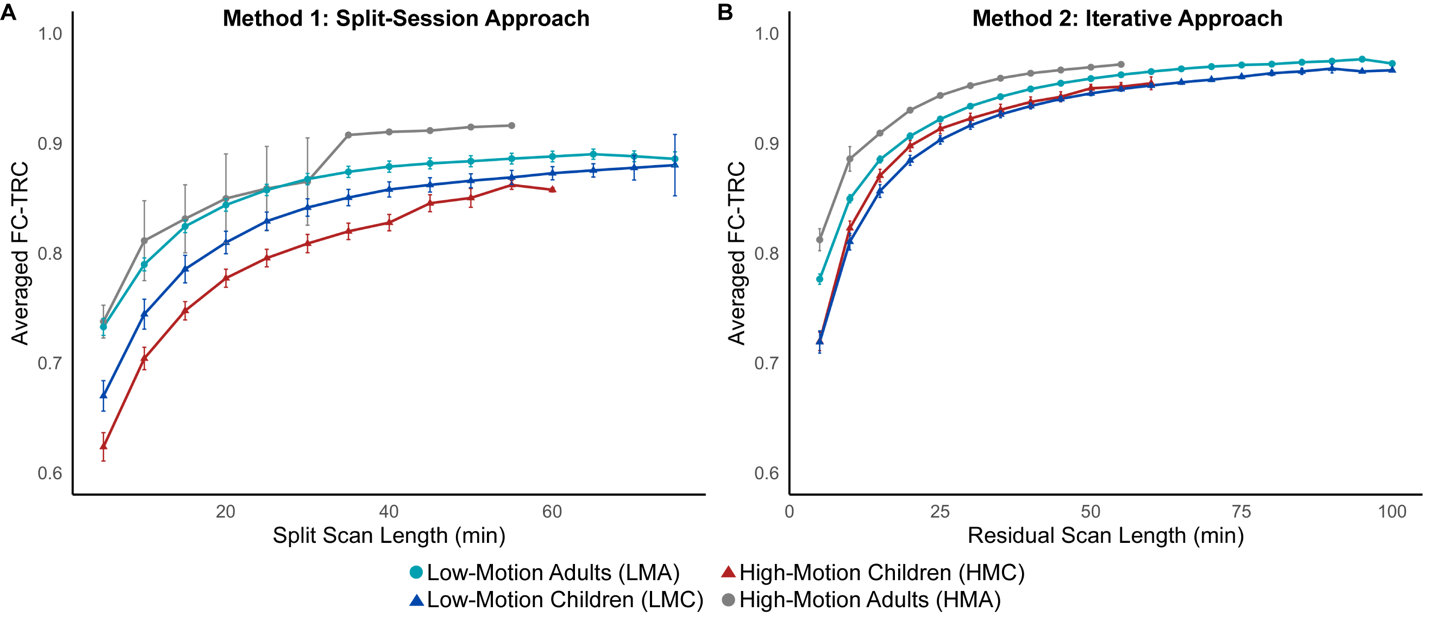

**Supplemental Figure 5: Group averaged FC-TRC-by-scan duration curves combined across all viewing conditions.** Averaging across participants in each motion group, both the split-session approach (A) and iterative approach (B) revealed differences between the low-motion adults (LMA, light blue circles) and both child groups. A) The split-session approach indicated that the low-motion children (LMC, dark blue triangles) attained higher FC-TRC values compared to the high-motion children (HMC, red triangles). Compared to (A), the iterative approach in (B) had a smaller gap in FC-TRC differences between both child groups and the low-motion adult group. The high-motion adult group (HMA, grey circles) included two participants that were excluded from further motion group analyses.

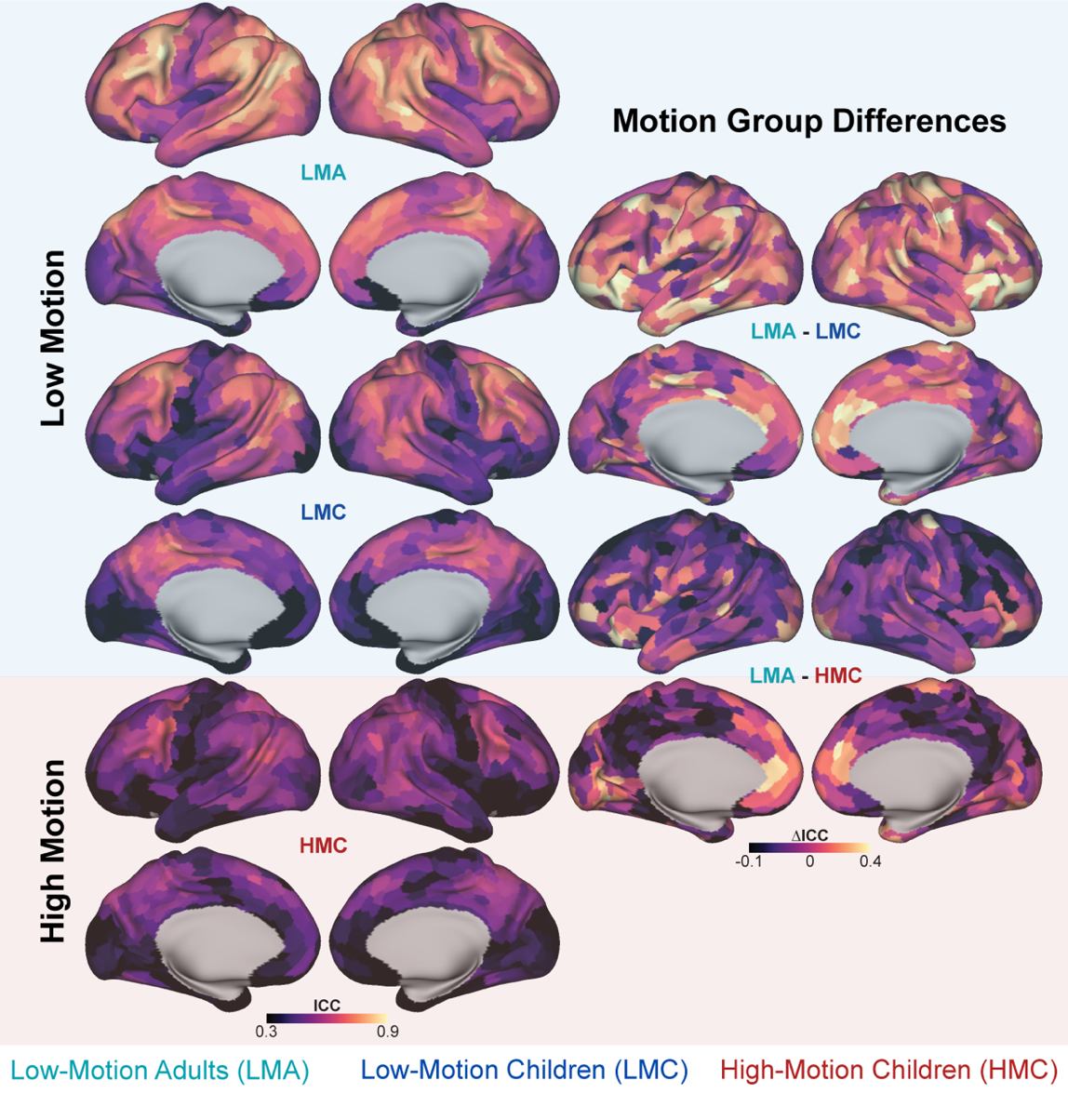

**Supplemental Figure 6: Regional differences in Intraclass Correlation Coefficient (ICC) reliability across motion groups under passive viewing conditions.** The low-motion adult (LMA) group exhibited greater overall ICC reliability with difference maps between groups shown in the right column. The largest difference observed was between the low-motion adults (LMA) and high-motion children (HMC) in frontal and temporal lobe regions. The Schaefer 1000 parcel 17-network atlas was used to parcellate brain regions.

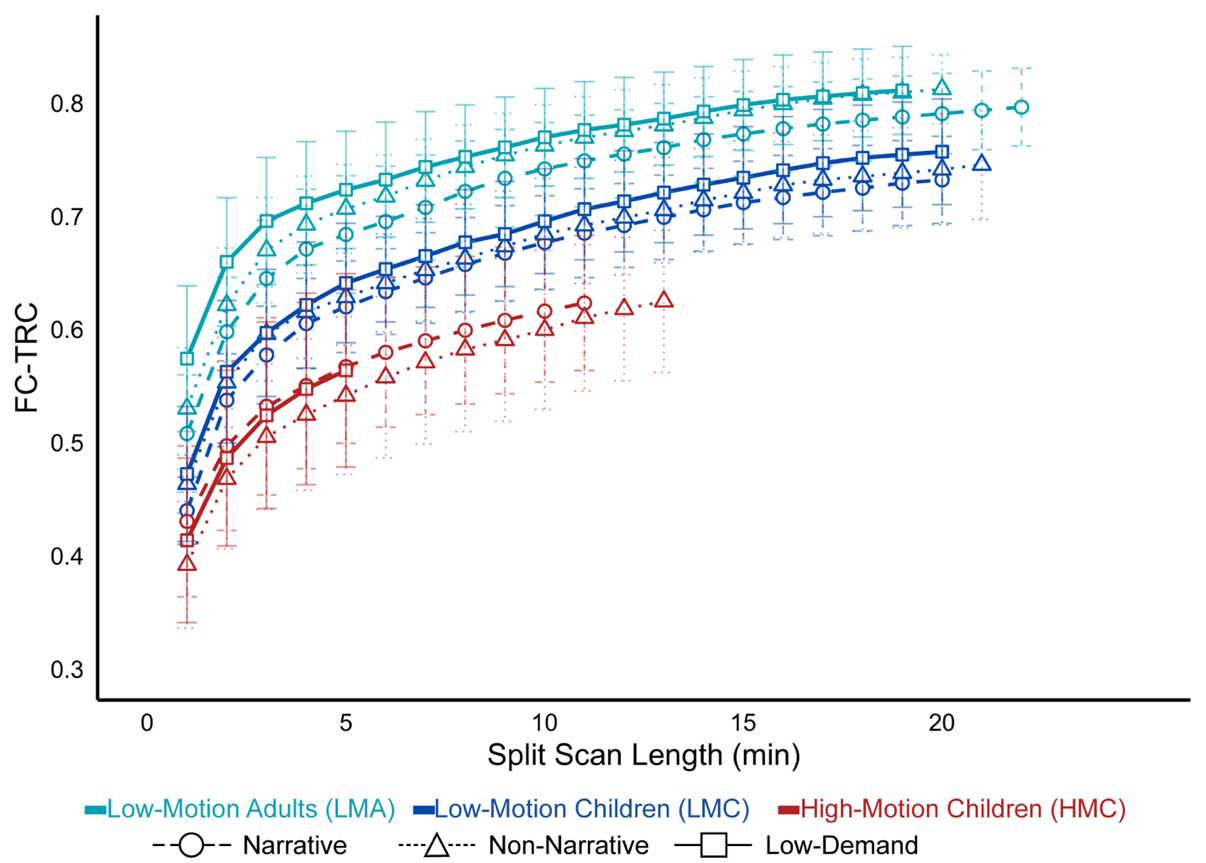

**Supplemental Figure 7: Test-retest correlations (FC-TRC) curves across each viewing condition.** The low-motion adults (LMA, light blue) showed higher FC-TRC values compared to both low-motion children (dark blue) and high-motion children (red) across all conditions. Across all motion groups, the low-demand condition, indicated by squares, achieved the highest FC-TRC, however, retained the least split-half data for the high-motion children (HMC) at 5 minutes.
